# *Pyricularia graminis-tritici* sp. nov., a new *Pyricularia* species causing wheat blast

**DOI:** 10.1101/051151

**Authors:** V.L. Castroagudín, S.I. Moreira, D.A.S. Pereira, S.S. Moreira, P.C. Brunner, J.L.N. Maciel, P.W. Crous, B.A. McDonald, E. Alves, P.C. Ceresini

## Abstract

Abstract *Pyricularia oryzae* is a species complex that causes blast disease on more than 50 species of poaceous plants. *Pyricularia oryzae* has a worldwide distribution as a rice (*Oryza*) pathogen and in the last 30 years emerged as an important wheat (*Triticum*) pathogen in southern Brazil. We conducted phylogenetic analyses using 10 housekeeping loci for 128 isolates of *P. oryzae* sampled from sympatric populations of grasses growing in or near wheat fields. Phylogenetic analyses grouped the isolates into three major clades. Clade 1 comprised isolates associated only with rice and corresponds to the previously described rice blast pathogen *P. oryzae* pathotype *Oryza* (PoO). Clade 2 comprised isolates associated almost exclusively with wheat and corresponds to the previously described wheat blast pathogen *P. oryzae* pathotype *Triticum* (PoT). Clade 3 contained isolates obtained from wheat as well as other *Poaceae* hosts. We found that Clade 3 is distinct from *P. oryzae* and represents a new species, *Pyricularia graminis-tritici*, (Pgt). No morphological differences were observed among these species, but a distinctive pathogenicity spectrum was observed. Pgt and PoT were pathogenic and highly aggressive on *Triticum aestivum* (wheat), *Hordeum vulgare* (barley), *Urochloa brizantha* (signal grass) and *Avena sativa* (oats). PoO was highly virulent on the original rice host (*Oryza sativa*), and also on wheat, barley, and oats, but not on signal grass. We conclude that blast disease on wheat and its associated *Poaceae* hosts in Brazil is caused by multiple *Pyricularia* species. *Pyricularia graminis-tritici* was recently found causing wheat blast in Bangladesh. This indicates that *P. graminis-tritici* represents a serious threat to wheat cultivation globally.

## INTRODUCTION

*Pyricularia oryzae* is a species complex (Couch & Kohn 2002) that causes blast disease on more than 50 species of poaceous plants, including important crops such as rice, wheat, barley, millet and oats (Urashima & Kato 1998, Couch & Kohn 2002, Takabayashi et al. 2002, Murakami et al. 2003, Couch et al. 2005). On the basis of host specificity, mating ability, and genetic relatedness, *P. oryzae* isolates were classified into several subgroups with restricted host ranges, including: the *Oryza* pathotype, pathogenic on rice (*Oryza sativa*); the *Setaria* pathotype, pathogenic on foxtail millet (*Setaria italica*); the *Panicum* pathotype, pathogenic on common millet (*Panicum miliaceum*); the *Eleusine* pathotype, pathogenic on finger millet (*Eleusine coracana*); the *Triticum* pathotype, pathogenic on wheat (*Triticum aestivum*); the *Avena* pathotype, pathogenic on oats (*Avena sativa*); and the *Lolium* pathotype, pathogenic on perennial ryegrass (*Lolium perenne*) (Urashima et al. 1993, Kato et al. 2000, Tosa et al. 2004, Tosa & Chuma 2014). Kato and collaborators (Kato et al. 2000) reported that isolates of *P. oryzae* recovered from *Eleusine, Panicum, Oryza, Setaria, and Triticum* spp. form a highly related group that is partially inter-fertile with the *Oryza* subgroup (i.e. the rice blast pathogen). In addition, the *Oryza* and *Setaria* pathotypes contain physiological races that show distinct patterns of virulence on cultivars within their host species (Tosa & Chuma 2014). Both host species-specificity and cultivar-specificity can be governed by gene-for-gene interactions (Silue et al. 1992, Takabayashi et al. 2002, Tosa et al. 2006, Valent & Khang 2010).

The *P. oryzae* pathotype *Triticum* is considered the causal agent of wheat blast in South America and has also been associated with blast disease on barley, rye, triticale, and signal grass (*Urochloa* sp.) in central-western and southern Brazil (Lima & Minella 2003, Verzignassi et al. 2012). Wheat blast was first reported in Paraná State, Brazil in 1985 (Igarashi et al. 1986, Anjos et al. 1996). Due to the lack of resistant cultivars and effective fungicides for disease management, wheat blast is widely distributed across all the wheat-cropping areas in Brazil, causing crop losses from 40-100% (Silva et al. 2009, Maciel 2011, Castroagudín et al. 2015). Wheat blast also occurs in Bolivia, Argentina and Paraguay (Duveiller et al. 2010). The disease was not found outside South America (Maciel 2011) until a recent outbreak reported in Bangladesh (Callaway 2016), though wheat blast is considered a major quarantine disease and a threat to wheat crops in the United States (Duveiller et al. 2007, Kohli et al. 2011).

As wheat blast emerged in an area of southern Brazil where rice blast is prevalent, it was originally proposed that the rice pathogen had evolved to parasitize wheat (Igarashi et al. 1986). Urashima et al. (1993) provided evidence based on pathogenicity, reproductive isolation and genetic data that indicated the existence of two distinct groups of *P. oryzae* causing wheat blast in Brazil, one that infects rice and wheat and one that infects only wheat. In that study, wheat-derived isolates were reported to infect grass plants from six different tribes within *Poaceae*. In addition, crosses of wheat-derived isolates with strains from *Eleusine coracana, Urochloa plantaginea (*ex *Brachiaria plantaginea*), and *Setaria italica* produced mature perithecia with viable ascospores, i.e. evidence of fertile crosses (Urashima et al. 1993). On the contrary, progeny from the crosses between wheat-and rice-derived isolates were infertile (Urashima et al. 1993). In the same study, crosses between wheat-derived isolates and isolates obtained from *Cenchrus echinatus, Setaria geniculata, Echinocloa colonum* produced no perithecia (Urashima et al. 1993). The work of Urashima and his collaborators indicated that two distinct pyricularia-like pathogens cause wheat blast disease in Brazil. However, it is not clear whether a population of *P. oryzae* able to infect both rice and wheat coexists with a population that infects only wheat.

Several studies suggested that the wheat-adapted *P. oryzae* population was derived *de novo* from a non-rice host. DNA fingerprinting with the repetitive DNA probes MGR563 and MGR586 found a high level of differentiation between *P. oryzae* pathotype *Oryza* (PoO) and *P. oryzae* pathotype *Triticum* (PoT) from Brazil (Farman 2002). In fact, the fingerprints from wheat-derived isolates resembled those from isolates non-pathogenic to rice (Hamer 1991, Valent & Chumley 1991, Urashima et al. 1999, Farman 2002). Maciel et al. (2014) showed that the Brazilian wheat-adapted population of *P. oryzae* was highly differentiated (F_CT_ = 0.896, *P* ≤ 0.001) from the local rice-adapted population. Analyses of the current pathotype diversity of *P. oryzae* showed that none of the 69 wheat-derived isolates were able to infect rice (Maciel et al. 2014).

Phylogenetic analyses demonstrated that *Pyricularia* is a species-rich genus in which different species evolved through repeated radiation events from a common ancestor (Hirata et al. 2007, Choi et al. 2013, Klaubauf et al. 2014). Multi-locus phylogenetic analyses revealed that *P. oryzae* and *P. grisea* are independent phylogenetic species (Taylor et al. 2000, Couch & Kohn 2002) and showed that the contemporary rice-infecting pathogen (*P. oryzae* pathotype *Oryza*) originated via a host shift from millet onto rice ~7,000 years ago during rice domestication in China (Couch et al. 2005). More recent phylogenetic analyses combined pre-existing biological and morphological data to re-examine the relationships among pyricularia-like species. These comprehensive studies favoured the classification of new cryptic species that were recently identified within *Pyricularia* and other relevant changes within the order *Magnaporthales* (Hirata et al. 2007, Choi et al. 2013, Luo & Zhang 2013, Klaubauf et al. 2014, Murata et al. 2014). Most relevant for agricultural scientists is that despite the extensively reported differentiation between *P. oryzae* pathotypes *Oryzae* and *Triticum*, these two pathotypes have been kept under the same species name *P. oryzae*. Therefore, we sought to determine whether the pathotypes *Oryza* and *Triticum* of *P. oryzae* are distinct species that should be given different names. We conducted phylogenetic analyses based on 10 housekeeping genes using sympatric populations of *Pyricularia* sampled from rice, wheat and other poaceous hosts in Brazil. We also conducted cultural, morphological, and pathogenic characterisation of the *Pyricularia* isolates to provide a complete description for each species. Our phylogenetic analyses revealed a new *Pyricularia* species causing blast on wheat and other poaceous hosts in Brazil. We name and describe *Pyricularia graminis-tritici* in this report.

## MATERIALS AND METHODS

### Fungal isolates and DNA extraction

A unique collection of 128 monoconidial isolates of *Pyricularia* spp. obtained in sympatry from the Brazilian wheat agro-ecosystem was analysed in this study (Table 1). *Pyricularia* spp. isolates were obtained from *Triticum aestivum* (*N* = 79), *Oryza sativa* (*N* = 23), *Avena sativa* (*N* = 5), *Cenchrus echinatus* (*N* = 3), *Cynodon* sp. (*N* = 1), *Digitaria sanguinalis* (*N* = 4), *Elionurus candidus* (*N* = 2), *Echinochloa crusgalli* (*N* = 1), *Eleusine indica* (*N* = 1), *Rhynchelytrum repens* (*N* = 3), and *Urochloa brizantha* (*N* = 6). Isolates recovered from wheat and other poaceous hosts located within or adjacent to sampled wheat plots were obtained from symptomatic leaf tissue in commercial wheat fields located in seven states in Brazil during 2012. A detailed description of wheat field sampling strategies was provided earlier (Castroagudín et al. 2015). The rice-derived isolates of *P. oryzae* were recovered from rice leaves, necks and panicles exhibiting typical rice blast symptoms, comprising a representative group including all races of *P. oryzae* subgroup *Oryza* prevalent in the major Brazilian rice growing areas (Maciel et al. 2014). The rice-derived isolates were provided by EMBRAPA-Rice and Beans, Santo Antônio de Goiás, Goiás, Brazil. The isolate collection is maintained at the Laboratory of Phytopathology, UNESP-DEFERS Campus Ilha Solteira, São Paulo, Brazil. A duplicate of the collection is hosted at the Laboratory of Phytopathology, EMBRAPA-Wheat, Passo Fundo, Brazil. Specimens were deposited at Culture Collection Mycobank Prof.

**Table 1.**
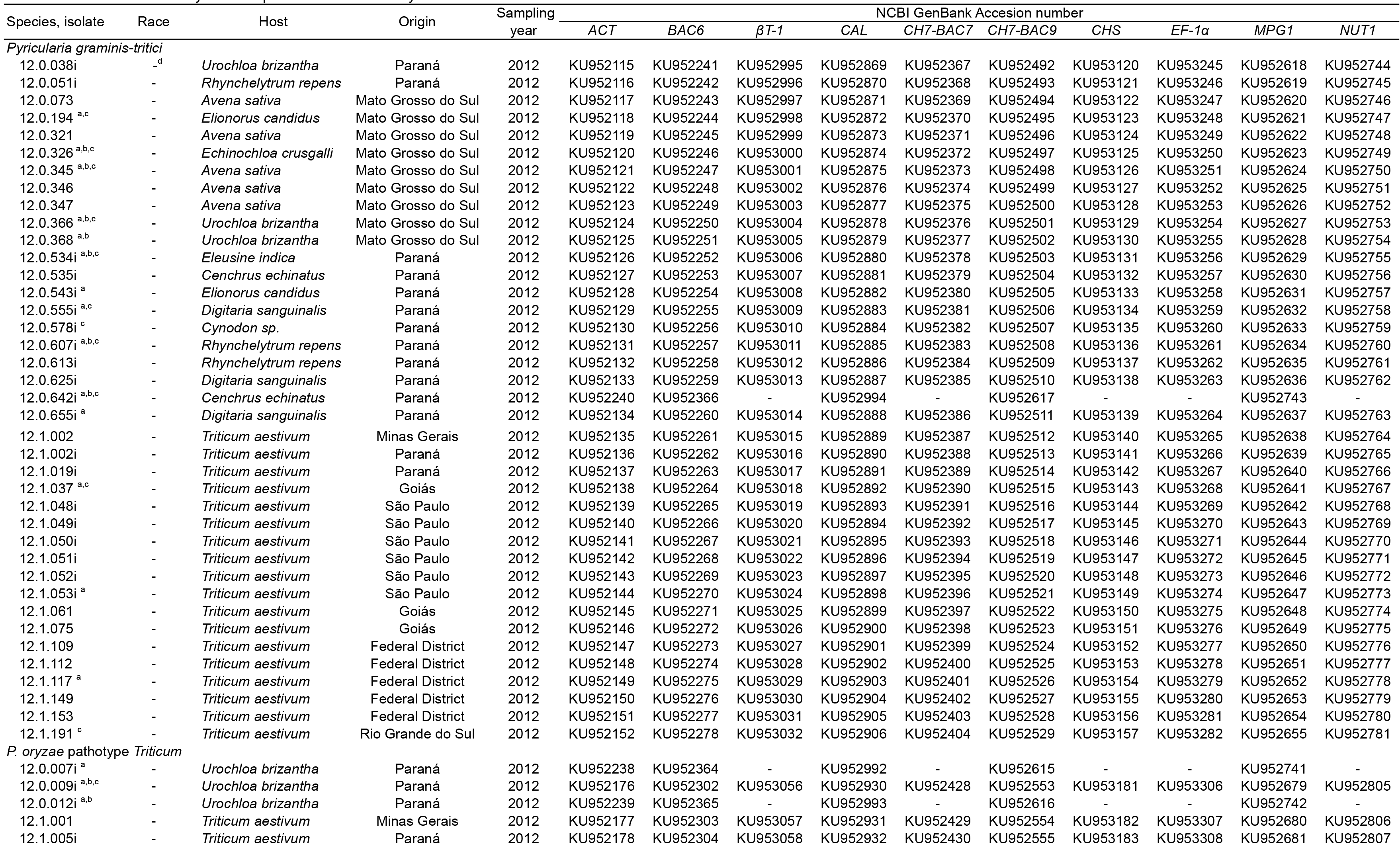
Details of isolates of *Pyricularia* spp. used in this study and NCBI Accession numbers

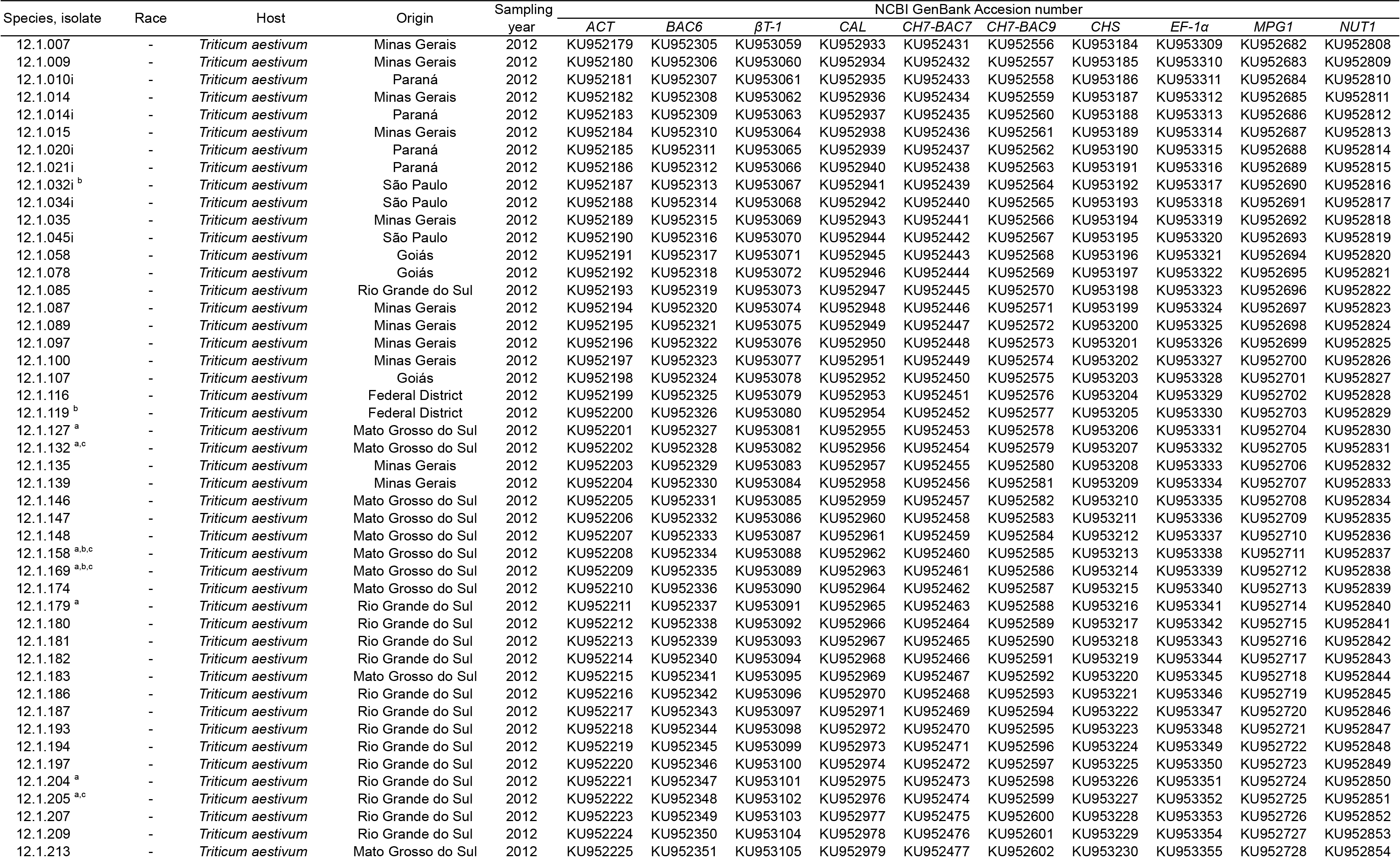

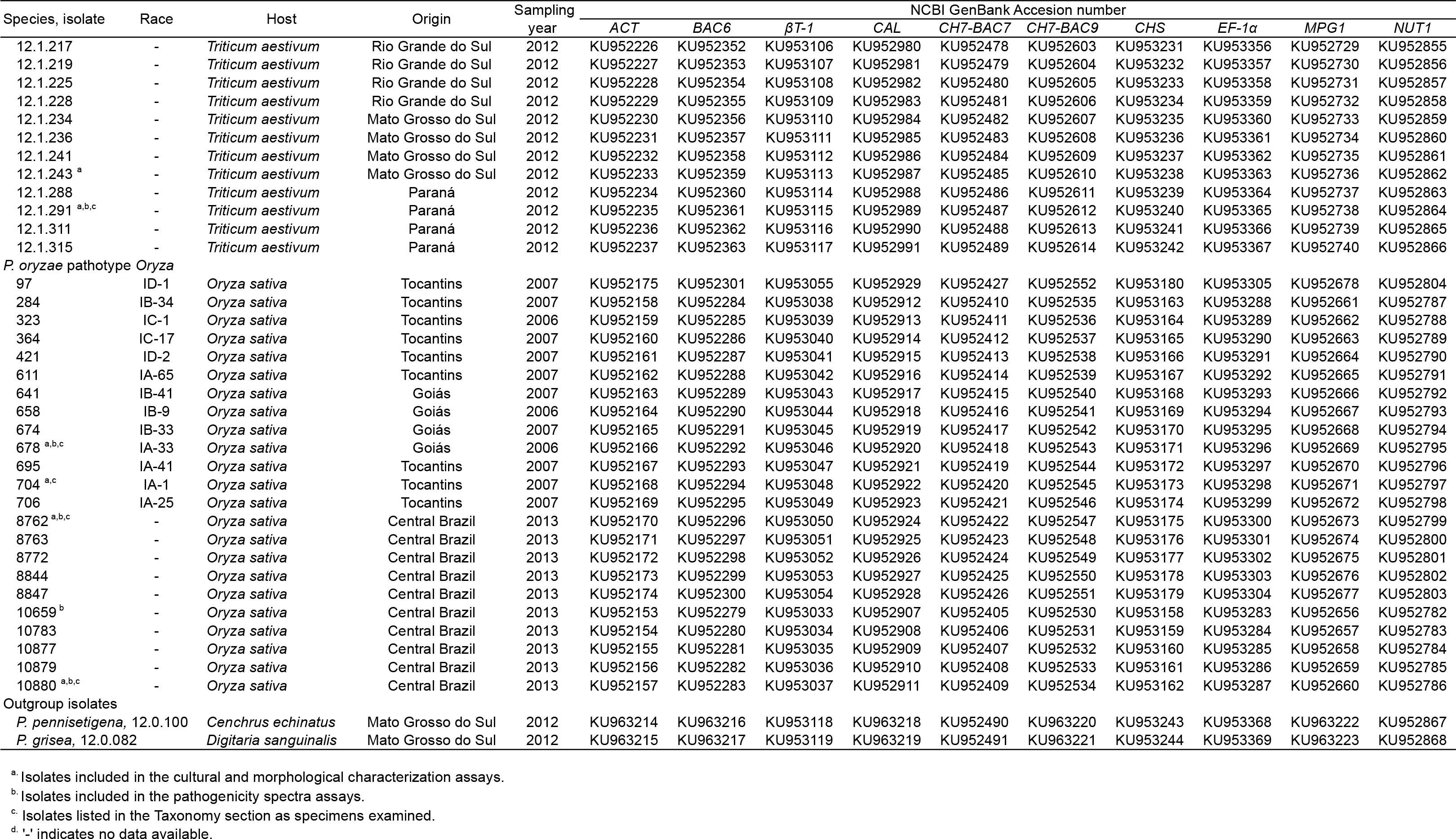

Maria Auxiliadora Cavalcanti, Federal University of Pernambuco, Recife, Brazil; and at the Coleção de Culturas da Microbiologia Agrícola (Agriculture Microbiology Culture Collection) of the Federal University of Lavras, Lavras, Minas Gerais, Brazil. Holotype specimen was deposited at INCT-HISA Herbário Virtual da Flora e dos Fungos at UNESP – Campus Ilha Solteira (Virtual Herbarium of Flora and Fungi, University of São Paulo State – Campus Ilha Solteira, Ilha Solteira, São Paulo, Brazil).

### DNA extraction, amplification and sequencing

Genomic DNA was extracted from freeze-dried mycelia with the GenElute Plant Genomic DNA Miniprep Kit (Sigma-Aldrich, St. Louis, MO, USA), according to the specifications of the manufacturer. Partial sequences of 10 nuclear housekeeping loci previously used to characterise *Pyricularia* species (Carbone & Kohn 1999, Couch & Kohn 2002, Couch et al. 2005, Zhang & Zhao 2011) were included in the analyses. The loci amplified were: *ACT*(actin), *BAC6* (putative vacuolar import and degradation protein), *βT-1* (beta-tubulin), *CAL* (calmodulin), *CH7-BAC7* (hypothetical protein), *CH7-BAC9* (anonymous sequence), *CHS1* (chitin synthase 1), *EF-1α* (translation elongation factor 1-alpha), *MPG1* (hydrophobin), and *NUT1* (nitrogen regulatory protein 1). The loci were amplified using PCR cycling conditions described previously (Carbone & Kohn 1999, Couch et al. 2005). The PCR primers and the annealing temperatures used to amplify each locus are described in Table 2. The PCR products were purified and sequenced by Macrogen Inc. (Seoul, Korea) using the ABI Prism BigDye Terminator v.3.1 Cycle Sequencing Ready Reaction Kit in an ABI 3730xl automated sequencer (Applied Biosystems, Foster City, CA). Newly generated DNA sequences were deposited in NCBI's GenBank nucleotide database (Table 1).

**Table 2.**
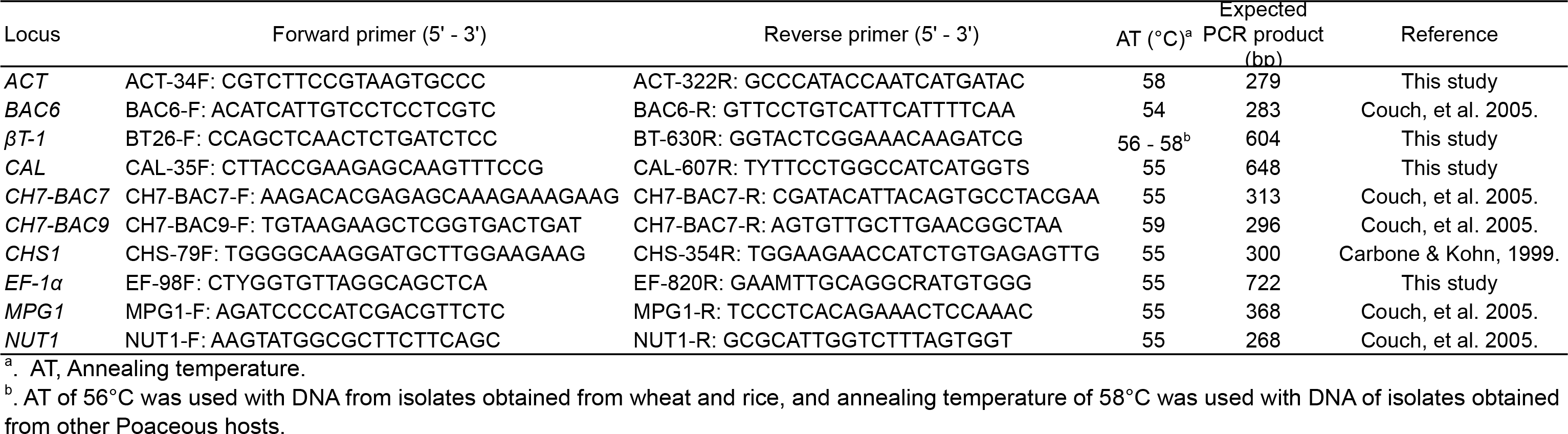
Primers used in this study

### Phylogenetic analyses

The complete set of sequence data was obtained from 125 isolates of *Pyricularia* spp., including two identified as *P. pennisetigena* (URM7372 = CLM3524, isolate 12.0.100) and *P. grisea* (URM7371 = CLM3525, isolate 12.0.082) from Brazil, which were used as outgroups. Sequence data from the 10 loci were assembled, aligned, and concatenated using Geneious R v. 9.0.5 (Biomatters, Auckland, New Zealand) for further phylogenetic analyses.

The phylogeny for the *Pyricularia* species was reconstructed through Bayesian inference using BEAST v 1.8.2 and in-files created with the help of BEAUti (Drummond et al. 2012). The 10-locus data set was partitioned and the best substitution model for each locus was determined using JModelTest2 (Darriba et al. 2012). Exploratory BEAST runs were conducted to determine the optimal clock-and tree-models. Model comparisons were based on the likelihoods using the Akaike information criterion (AICM) as implemented in the program Tracer v. 1.6 (Rambaut et al. 2014). The selected nucleotide substitution model was GTR for all loci, the strict clock model and the birth-death speciation process as the tree model.

Four independent final runs were conducted with MCMC length set to 10^8^ generations with sampling intervals every 1,000 generations. Runs were assessed for convergence and combined using LogCombiner v1.8.0, which is part of the BEAST package. Posterior sampled trees were extracted using TreeAnnotator v. 1.8.2. (Drummond et al. 2012) with the following parameters: burn-in 10%, 0.50 posterior probability limit, maximum clade credibility target tree type, and mean node height. The final tree was visualised with FigTree v. 1.4.2 (Institute of Evolutionary Biology, University of Edinburgh, http://tree.bio.ed.ac.uk/software/figtree). A phylogenetic tree was reconstructed for *MPG1* using the same settings as described for the combined tree. The resulting trees and respective alignments were deposited into TreeBASE (submission 19365). Based on the phylogenetic results, non-fixed and fixed nucleotide differences across all loci among the major clades were calculated using DnaSP (Librado & Rozas 2009).

### Cultural characterisation

To examine macroscopic features, a representative subgroup of 30 isolates (Table 1) were grown on Corn Meal Agar (CMA), Malt Extract Agar (MEA), Oatmeal Agar (OA), Potato Dextrose Agar (PDA), and Synthetic Nutrient-poor Agar (SNA). All media were prepared as previously described (Crous et al. 2009) and amended with streptomycin sulphate (INLAB, São Paulo, Brazil) 0.05 g/L, and chloramphenicol (INLAB, São Paulo, Brazil) 0.05 g/L.

Stored isolates were re-activated on PDA. For this assay, a 6-mm-diam disk of colonized PDA from a 7-d-old re-activated culture was transferred to the centre of a Petri plate containing one of the media described above. Colony diameter and cultural features were assessed after 7 d of incubation at 25 °C under a 12 h dark/12 h fluorescent light regime, following the procedures described by Klaubauf et al. (2014). Three replicates were made for each isolate and the assay was conducted twice. For colony descriptions, isolates were grouped according to their clustering in the phylogenetic analysis. A general description representing the colony morphology of each group of isolates was recorded. In addition, one isolate from each group was chosen as representative of the group.

### Morphological characterisation

The same subgroup of 30 isolates selected for the description of colony morphology was examined using bright field and electron microscopy to characterise fungal structures. Isolates were re-activated on CMA and incubated for 7 d at 25 °C in darkness. They were subsequently transferred to SNA with sterile barley seeds to induce sporulation and incubated for 3 wk at 25 °C under a 12 h dark/12 h fluorescent light regime. Samples were prepared following methods described previously (Bozzola & Russell 1999).

Observations were made with a Nikon SMZ25 stereo-microscope, and with a Zeiss Axio Imager 2 light microscope using differential interference contrast (DIC) illumination and a Nikon DS-Ri2 camera and software. The bright field images were taken with a Nikon SMZ1500 stereoscope microscope using NIS Elements D 3.2 software. Scanning electron microscope (SEM) images and measurements were acquired on a Zeiss LEOEVO 40 microscope using SmartSem Zeiss software (Oberkochen, Germany) operating at 10 kV and 10 to 30 mm work distance. When possible, biometric data were obtained from 30 observations per fungal structure per isolate. The photo plates were created on Corel Draw X7 software (Corel Corporation, Ottawa, Canada).

### Pathogenicity spectrum

A subgroup of 18 isolates was tested for pathogenicity spectra in greenhouse assays on barley (*Hordeum vulgare*) cvs. BRS Korbel, signal grass (*Urochloa brizantha, ex Brachiaria brizantha*) cvs. Piata and Marandu, oats (*Avena sativa*) cvs. EMBRAPA 29 and IAPAR 61, rice (*Oryza sativa*) cv. IRGA 409, and wheat cv. Anahuac 75. Seeds of the different hosts were planted in 10-cm-diam plastic pots filled with Tropstrato HT potting mix (Vida Verde, Mogi Mirim, São Paulo, Brazil). Fifteen seeds were planted per pot. Fifteen d after seedling emergence, pots were thinned to eight seedlings per pot for barley, signal grass, oats, and rice; and to five seedlings per pot for wheat. Pots were kept in the greenhouse under natural conditions until inoculation and watered daily from the top. Plants were fertilised with NPK 10:10:10 granular fertiliser (N:P_2_O_5_:K_2_O, Vida Verde, Mogi Mirim, São Paulo, Brazil). A forty gram dose of NPK granular fertiliser was sprinkled across every 100 pots 1 d after emergence. Fertilisation was repeated every 15 d until inoculation. In addition, rice plants were fertilised with a solution of 4 g/L FeSO_4_7H_2_O (Dinâmica, Diadema, São Paulo, Brazil) once after emergence, with 1 L of solution applied to every 100 pots.

Isolates were recovered from long-term storage and re-activated on PDA plates and then transferred either to OA plates (rice-derived isolates) or PDA plates (wheat and other isolates originating from poaceous hosts). Fifteen plates were prepared for each isolate. Plates were incubated for 15 d at 25 °C under a 12 h dark/12 h fluorescent light regime. Mycelium was gently scraped and washed with 3-5 mL of sterile distilled water amended with Tween 80 (two drops/L) to release the spores. Conidia concentration was quantified using a Neubauer counting chamber and adjusted to 1 × 10^5^ spores/mL for inoculation.

Pathogenicity assays were conducted on seedlings, 1-mo-old plants at growth stage 14 (Zadocks et al. 1974) on all hosts, and on immature heads of 2-mo-old wheat plants at the beginning of anthesis in growth stage 60 (Zadocks et al. 1974). Spore suspensions (1 × 10^5^ spores/mL) were uniformly applied either onto the adaxial leaf surfaces or onto wheat heads until runoff. Fifty millilitres of spore suspension was used for every 20 inoculated pots.

Inoculated pots were placed onto plastic trays and incubated in a plant growth chamber for 7 d at 26 °C (barley, oats, rice, and wheat) or 30 °C (signal grass). Plants were kept in the dark for the first 24 h, followed by a 12 h dark/12 h fluorescent light regime. Plants were watered every other day from the bottom to avoid cross-contamination. Humidifiers were used to insure that relative humidity would stay above 85% within the chamber during the entire experiment.

Temperature and relative humidity were recorded in the chamber using an ITLOG80 Datalogger (Instrutemp, Belenzinho, São Paulo, Brazil). As negative controls, five pots of each host were mock-inoculated with sterile deionised water amended with Tween 80 at two drops/L in each experimental replication.

Plants were examined for lesions 7 d after inoculation. For the seedling inoculation tests, the disease severity index was calculated using an ordinal scale from 0 to 5 as previously described (Urashima et al. 2005). The disease severity index was scored as follows: lesion type 0 = no visible reaction; 1 = minute, pinhead-sized spots; 2 = small brown to dark brown lesions with no distinguishable centres; 3 = small eyespot shaped lesions with grey centres; 4 = typical elliptical blast lesions with grey centres; 5 = completely dead plant. Index values 0, 1, and 2 were considered non-compatible and index values 3, 4 and 5 were considered compatible. When different types of lesions were found on a single leaf, the most abundant lesions were considered.

Disease severity on wheat heads was assessed following the procedure described by Maciel et al. (2014), calculating the percentage of each wheat head affected by blast using Assess v. 2.0 image analysis software (APS, St. Paul, Minnesota). Wheat head tissue was considered affected by blast when it was chlorotic and/or it was covered with pathogen spores. For each head, a picture from each side of the head was taken, and the percentage of affected area in the two pictures was averaged. Seedling and head inoculation experiments were conducted using a one-factor completely randomized unbalanced design. Five pots containing five (wheat) or eight (barley, signal grass, oats, and rice) plants in the seedling tests, or five non-detached heads in the wheat-head tests were inoculated with each of the 18 isolates. The seeding inoculation experiments were conducted twice. The head inoculation experiment was conducted six times, but only two randomly chosen replicates were used for further statistical analyses. For statistical analyses, isolates were grouped according to their phylogenetic clustering (i.e. based on the species clades identified using the 10 loci sequences).

Analyses of variance (ANOVA) were performed to evaluate the effects of experiment's replicates, *Pyricularia* species, and their interactions in the different inoculation tests. Analyses were performed independently for each host species. For non-parametric data (seedlings inoculation tests) ANOVAs were conducted using the PROC NPAR1WAY procedure computed with the Wilcoxon rank-sum test and by using Monte Carlo estimations for the exact *p*-values (*P*) with the EXACT/ MC statement, at α = 0.01. A Dunn all Pairs for Joint Ranks test was used for non-parametric means comparisons. In the seedlings inoculation experiment, replicates were not significantly different (exact *P* ≥ 0.05), thus the two replicates were combined for these analyses. For parametric data (wheat heads inoculation tests) ANOVAs were conducted with the PROC GLM procedure, considering species as fixed factors and isolates as random factors nested inside species factors. Fisher's protected Least Significant Difference (LSD) test was used for comparison of disease severity means for species, at α = 0.05. Since the experiment was unbalanced, the harmonic cell size was used to calculate the average LSD. The interaction between species and experiment replicate was statistically significant (*P* = 0.02), therefore the two replicates of the experiment were analysed independently.All statistical analyses were performed with Statistical Analysis System program, v. 9.4 (SAS Institute, Cary, North Carolina)

## RESULTS

### Phylogenetic analyses

The final alignment for partial sequences of the 10 genes had a total length of 3 381 bases (3 301 un-gapped bases) from 125 isolates, including sequences retrieved from Brazilian isolates of *P. grisea* and *P. pennisetigena* used as outgroups. A total of 471 polymorphic sites were found, equivalent to 14.3% of the un-gapped alignment total length, and 168 of these sites (5.1%) were phylogenetically informative (Table 3). This resulted in 109 multilocus haplotypes, i.e. 87.2% of isolates had a unique multilocus haplotype.

**Table 3.** Number of polymorphic sites in ten loci across Pyricularia species examined in this study

| Locus | Alignment length (bp) | Un-gapped sequence mean length (bp) | Polymorphic sites<sup>a</sup> |  |
| --- | --- | --- | --- | --- |
|  |  |  | including outgroups<sup>b</sup> | excluding outgroups<sup>c</sup> |
| *ACT* | 184 | 179 | 16 (2)<sup>d</sup> | 0 (0) |
| *BAC6* | 254 | 253 | 18 (0) | 0 (0) |
| *βT-1* | 501 | 500 | 28 (9) | 19 (9) |
| *CAL* | 524 | 520 | 92 (33) | 12 (5) |
| *CH7-BAC7* | 285 | 285 | 54 (34) | 54 (34) |
| *CH7-BAC9* | 293 | 268 | 40 (20) | 38 (20) |
| *CHS* | 229 | 224 | 78 (8) | 26 (2) |
| *EF-1α* | 658 | 643 | 83 (31) | 66 (30) |
| *MPG1* | 229 | 205 | 55 (26) | 22 (16) |
| *NUT1* | 224 | 224 | 7 (5) | 5 (4) |
| Total | 3381 | 3301 | 471 (168) | 242 (120) |

<sup>a</sup>. Sequences of isolates 12.0.100 (*P. pennisetigena*, URM7372) and 12.0.082 (*P. grisea*, URM7371) were used as outgroups.
<sup>b</sup>. *N* = 125
<sup>c</sup>. *N* = 123
<sup>d</sup>. The number of phylogenetically informative sites is indicated between parenthesis.

The Bayesian analysis grouped the isolates into three major phylogenetic clades (Fig. 1). Clade 1 (Bayesian posterior probability, BPP = 1) comprised isolates exclusively associated with rice and corresponds to the previously described *P. oryzae* pathotype *Oryza* (PoO). Clade 2 (BPP = 0.99) comprised isolates almost exclusively associated with wheat. A single isolate (12.0.009i) collected from signal grass plants invading a wheat field in Paraná state also clustered within this clade. This clade corresponds to the previously described *P. oryzae* pathotype *Triticum* (PoT). Clade 3 (BPP = 0.99) contained isolates obtained from wheat as well as other *Poaceae* hosts. Based on the combined evidence presented in this study, we propose that this clade is distinct from *P. oryzae* and represents a new species, *Pyricularia graminis-tritici* (Pgt).

**Fig. 1.**
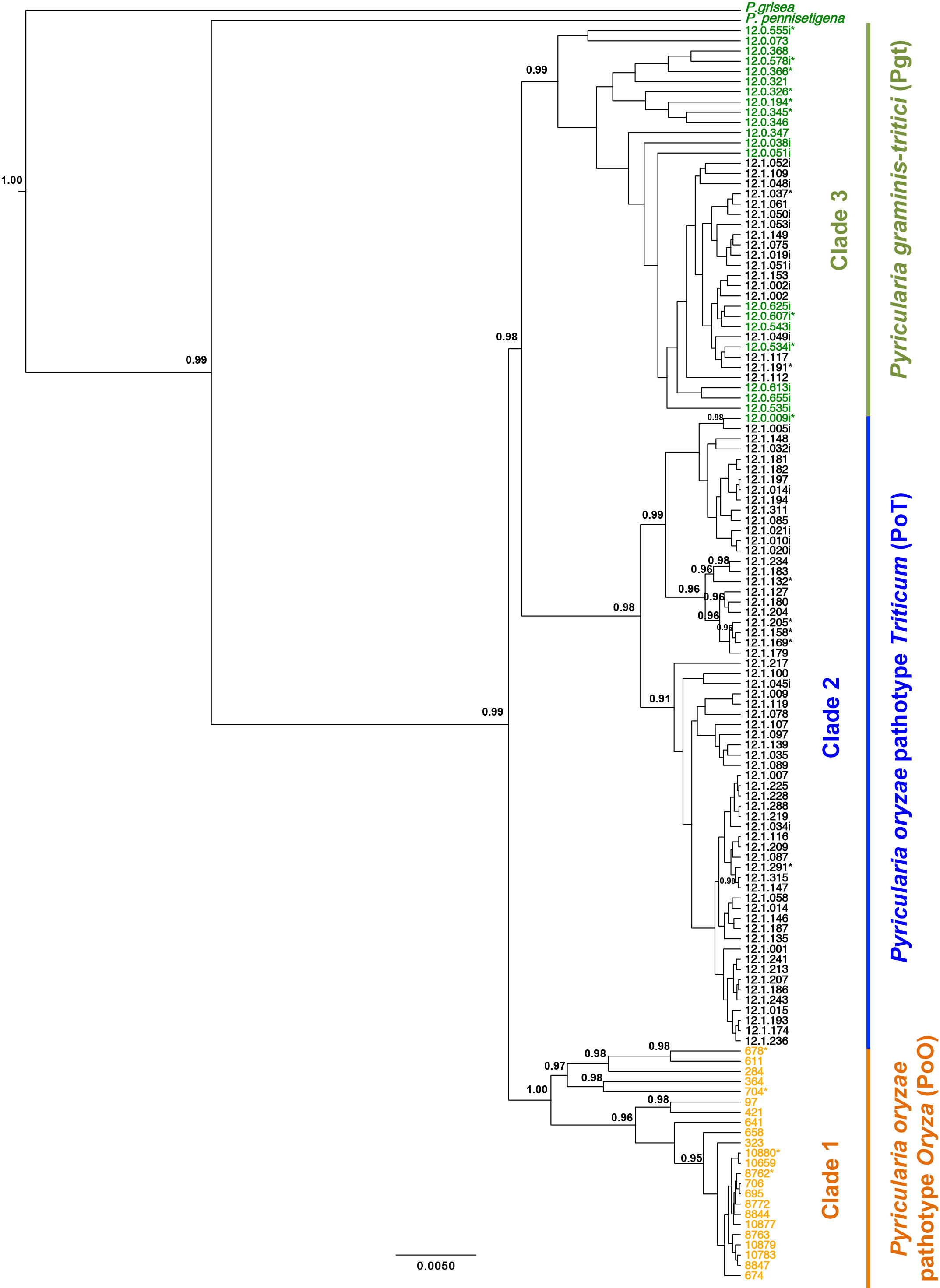
Phylogeny inferred by Bayesian Inference from the combined sequences of 10 partial loci (actin, *BAC6*, β-tubulin, calmodulin, *CH7-BAC7, CH7-BAC9*, chitin synthase 1, translation elongation factor 1-α, *MPG1* hydrophobin, and nitrogen regulatory protein 1) from isolates of *Pyricularia* spp. The 50% majority-rule consensus tree is shown. The numbers above the branches are the Bayesian posterior probabilities (BPP) for node support with BPP > 0.95. *Pyricularia grisea* and *P. pennisetigena* were used as out-groups. The original host of the isolate can be distinguished by the colour of the isolate number, black: wheat, green: other poaceous hosts and orange: rice. The asterisk (*) indicates the isolates listed in the Taxonomy section as specimens examined.

Non-fixed and fixed nucleotide differences among the three identified phylogenetic clades were examined for each locus, excluding the outgroups (Tables 3, 4). A total of 242 polymorphic sites were found, corresponding to 7.3% of the un-gapped alignment total length. Of those sites, 120 (3.6%) were phylogenetically informative. Four of the 10 loci (*βT-1, CH7-BAC9, EF-1α* and *MPG1*) showed a total of 18 (0.6%) fixed differences across the three clades (Tables 4, 5). *Pyricularia graminis-tritici* could be distinguished from PoT by 14 differences at *MPG1*. These fixed differences were at the following positions: 10 (C), 13-14 (TC), 20 (A), 22-25 (CCAG), 27 (C), 33—34 (CA), 41-42 (AG), and 87 (C). Likewise, Pgt could be distinguished from PoO by 18 fixed differences. These mutations are, one fixed difference at *βT-1*: 338 (A), one at *CH7-BAC9*: 20 (C), one at *EF-1α*: 325 (T) and 15 fixed differences at *MPG1*, as follows: 4 (T), 10 (C), 13-14 (TC), 20 (A), 22-25 (CCAG), 27 (C), 33-34 (CA), 41-42 (AG), and 87 (C). PoT was differentiated from PoO only by fixed differences: one difference at *CH7-BAC9*: 20 (C) and one at *EF-1α*: 325 (T), (Tables 4, 5).

**Table 4.** Number of fixed polymorphic sites in ten loci across *Pyricularia* species

| Species, clade | Locus | ACT | BAC6 | βT-1 | CAL | CH7-BAC7 | CH7-BAC9 | CHS | EF-1α | MPG1 | NUT1 | Total | %<sup>a</sup> |
| --- | --- | --- | --- | --- | --- | --- | --- | --- | --- | --- | --- | --- | --- |
|  | Alignment length (bp) | 184 | 254 | 501 | 524 | 285 | 293 | 229 | 658 | 229 | 224 | 3381 |  |
|  | Ungapped sequence mean length (bp) | 179 | 253 | 500 | 520 | 285 | 268 | 224 | 643 | 205 | 224 | 3301 |  |
| *P. graminis-tritici* vs. *P. oryzae* pathotype *Triticum* |  | 0 | 0 | 0 | 0 | 0 | 0 | 0 | 0 | 14 | 0 | 14 | 0.42 |
| *P. graminis-tritici* vs. *P. oryzae* pathotype *Oryza* |  | 0 | 0 | 1 | 0 | 0 | 1 | 0 | 1 | 15 | 0 | 18 | 0.55 |
| *P. oryzae* pathotype *Triticum* vs. *P. oryzae* pathotype *Oryza* |  | 0 | 0 | 0 | 0 | 0 | 1 | 0 | 1 | 0 | 0 | 2 | 0.06 |
| Total |  | 0 | 0 | 1 | 0 | 0 | 1 | 0 | 1 | 15 | 0 | 18 | 0.55 |

<sup>a</sup>. Percentage of fixed mutation with reference to the total number of 3301 nucleotides in the ungapped alignment.

**Table 5.** Fixed polymorphic sites in four loci across *Pyricularia* spp.

| | Locus | $\beta T-1$ | CH7-BAC9 | EF-1α | MPG1 | | | | | | | | | | | | | | | | |
| --- | --- | --- | --- | --- | --- | --- | --- | --- | --- | --- | --- | --- | --- | --- | --- | --- | --- | --- | --- | --- | --- |
| Species, clade | Alignment position | 776 | 1771 | 2597 | 2934 | 2940 | 2943 | 2944 | 2950 | 2952 | 2953 | 2954 | 2955 | 2957 | 2964 | 2965 | 2973 | 2974 | 3019 |  |  |
|  | Locus position | 338 | 20 | 325 | 4 | 10 | 13 | 14 | 20 | 22 | 23 | 24 | 25 | 27 | 33 | 34 | 41 | 42 | 87 |  |  |
| *Pyricularia graminis-tritici* |  | A | C | T | T | C | T | C | A | C | C | A | G | C | C | A | A | G | C |  |  |
| *P. oryzae* pathotype *Triticum* |  | A/C | C | T | T/C | T | C | G | C | T | T | C | - | T | T | C | - | - | A |  |  |
| *P. oryzae* pathotype *Oryza* |  | C | A | C | C | T | C | G | C | T | T | C | - | T | T | C | - | - | A |  |  |
| *P. pennisetigena* |  | A | C | C | T | A | A | T | T | A | T | C | A | T | T | C | - | G | A |  |  |
| *P. grisea* |  | C | C | C | A | T | T | T | C | A | T | G | G | C | C | G | A | - | A |  |  |

Sequences for only six genes were obtained for three isolates; therefore these isolates were not included in the phylogenetic analyses. However, by analysing variation in the diagnostic genes *CH7-BAC9* and *MPG1*, we were able to assign isolate 12.0.642i to Pgt, and isolates 12.0.007i and 12.0.012i to PoT.

### Cultural and morphological characterisation

For description of cultural and morphologic characteristics, *Pyricularia* isolates were grouped according to their phylogenetic placement, following the assignments *P. graminis-tritici* (Pgt), *P. oryzae* pathotype *Triticum* (PoT) and *P. oryzae* pathotype *Oryza* (PoO).

In general, similar colony morphologies were observed for isolates of Pgt, PoT, and PoO on the five media tested. No morphological differences were observed among the *Pyricularia* species. Cultural and morphological characteristics observed for *Pyricularia graminis-tritici* and *Pyricularia oryzae* pathotypes *Triticum* and *Oryza* (Fig. 3–5, a-j) are described in the Taxonomy section.

**Fig. 2.**
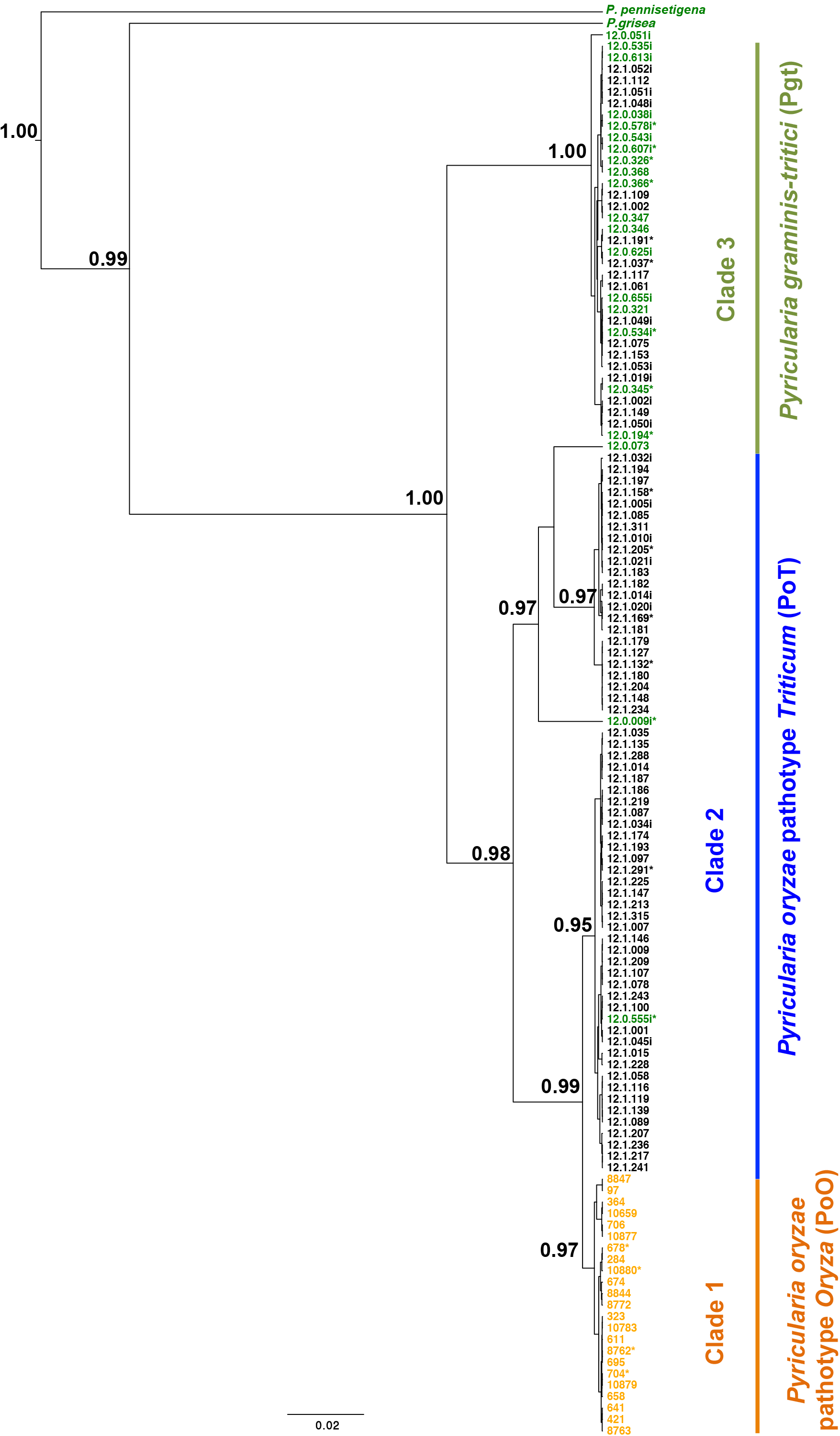
Phylogeny inferred by Bayesian Inference from the sequences *MPG1* hydrophobin locus from isolates of *Pyricularia* spp. The 50% majority-rule consensus tree is shown. The numbers above the branches are the Bayesian posterior probabilities (BPP) for node support with BPP > 0.95. *Pyricularia grisea* and *P. pennisetigena* were used as out-groups. The original host of the isolate can be distinguished by the colour of the isolate number, black: wheat, green: other poaceous hosts and orange: rice. The asterisk (*) indicates the isolates listed in the Taxonomy section as specimens examined.

**Fig. 3.**
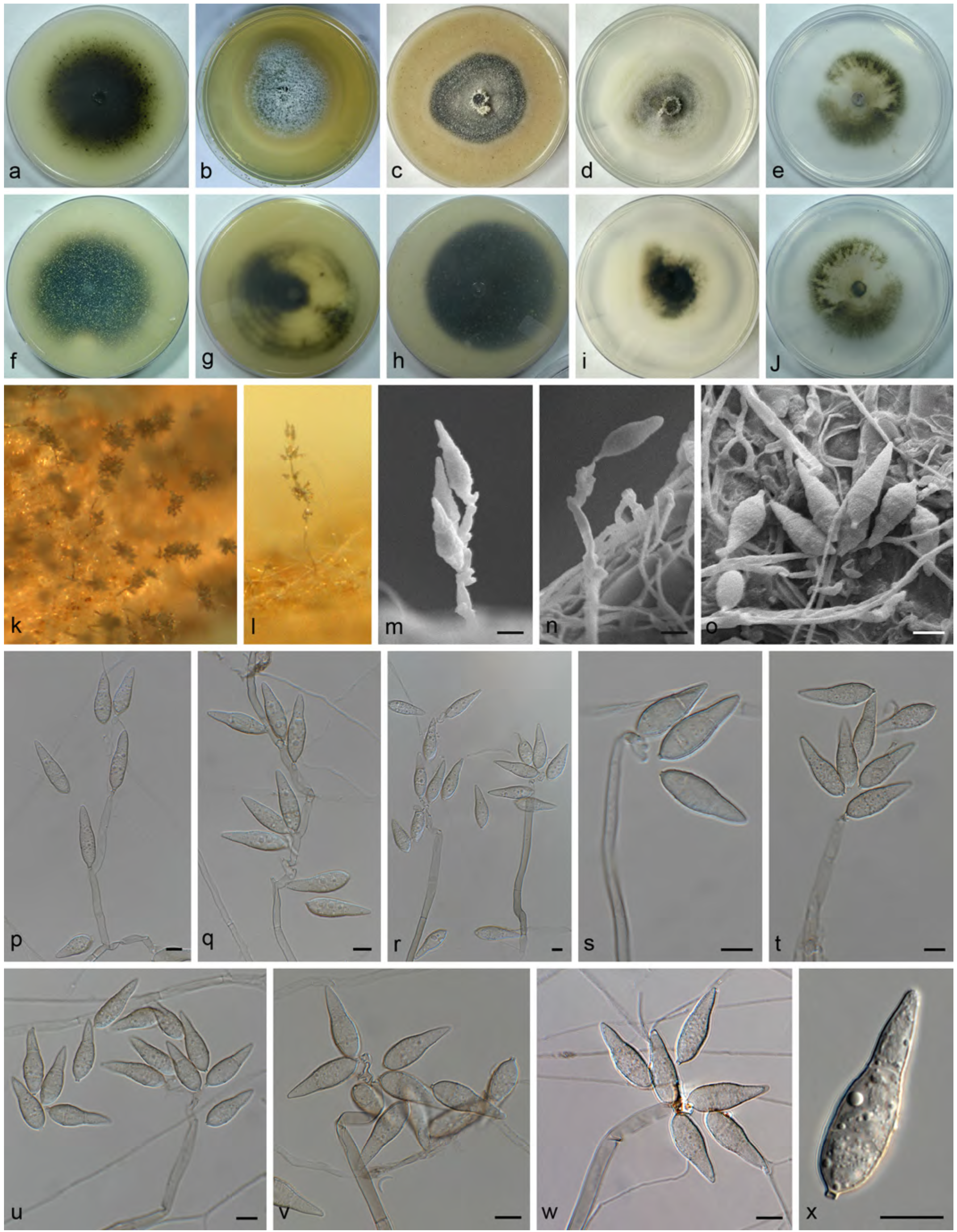
*Pyricularia graminis-tritici* a—j: cultures of isolate 12.1.037 grown for 7 d at 12 h photoperiod and 25°C in CMA (a, f), MEA (b, g), OA (c, h), PDA (d, i), and SNA (e, j) media; k—l, sporulation on SNA on sterile barley seeds; m-o, scanning electron micrographs of conidiophores and conidia; p—x, bright field microscopy images of conidiophores and conidia. – Scale bars = 10 μm.

### *Pathogenicity spectrum of pyricularia* spp. *on wheat, barley, signal grass, oats and rice*

The replicates of the seedlings inoculation tests were combined due to the lack of experiment effect (Table 6). *Pyricularia* species caused symptoms ranging from hypersensitive response lesions composed of diminutive, 1-mm-diam brown spots (DI = 1), to typical elliptical blast lesions with grey centres (> 5 mm diam), usually coalescing and causing plant death on all hosts (DI ≥ 3) (Kato et al. 2000, Cruz et al. 2016) (Fig. 6–8). This virulence variation was observed even among isolates of the same *Pyricularia* species and pathotypes, indicating the presence of host-physiological race interactions. For all tests, host seedlings or wheat heads used as negative controls showed no blast lesions on their leaves [mean disease index (DI) = 0.00].

**Fig. 4.**
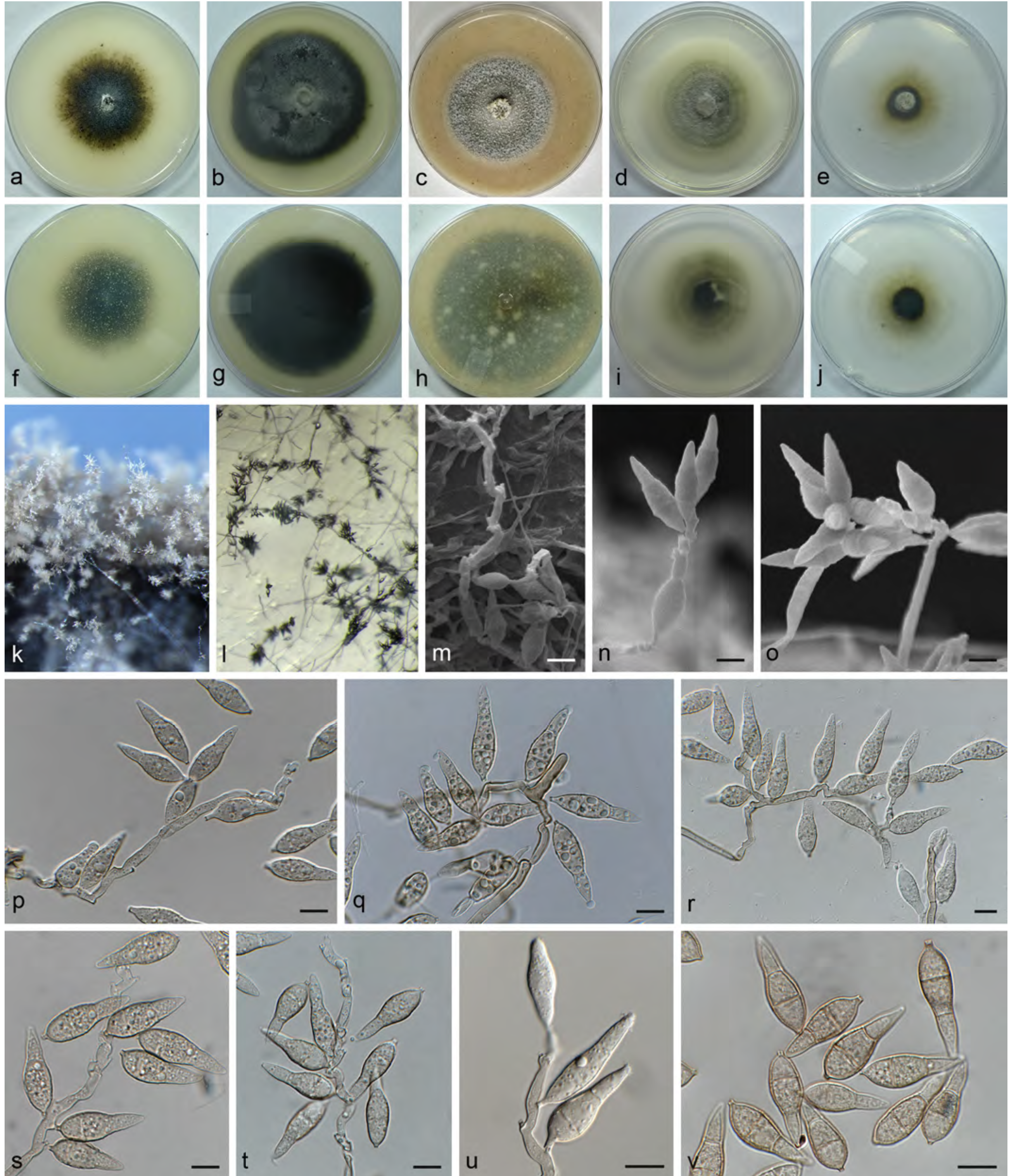
*Pyricularia oryzae* pathotype *Triticum*. a—j: cultures of isolate 12.1.291 grown for 7 d at 12 h photoperiod and 25°C in CMA (a, f), MEA (b, g), OA (c, h), PDA (d, i), and SNA (e, j) media; k—l, sporulation on SNA on sterile barley seeds; m—o, scanning electron micrographs of conidiophores and conidia; p—v, bright field microscopy images of conidiophores and conidia. – Scale bars = 10 μm.

**Fig. 5.**
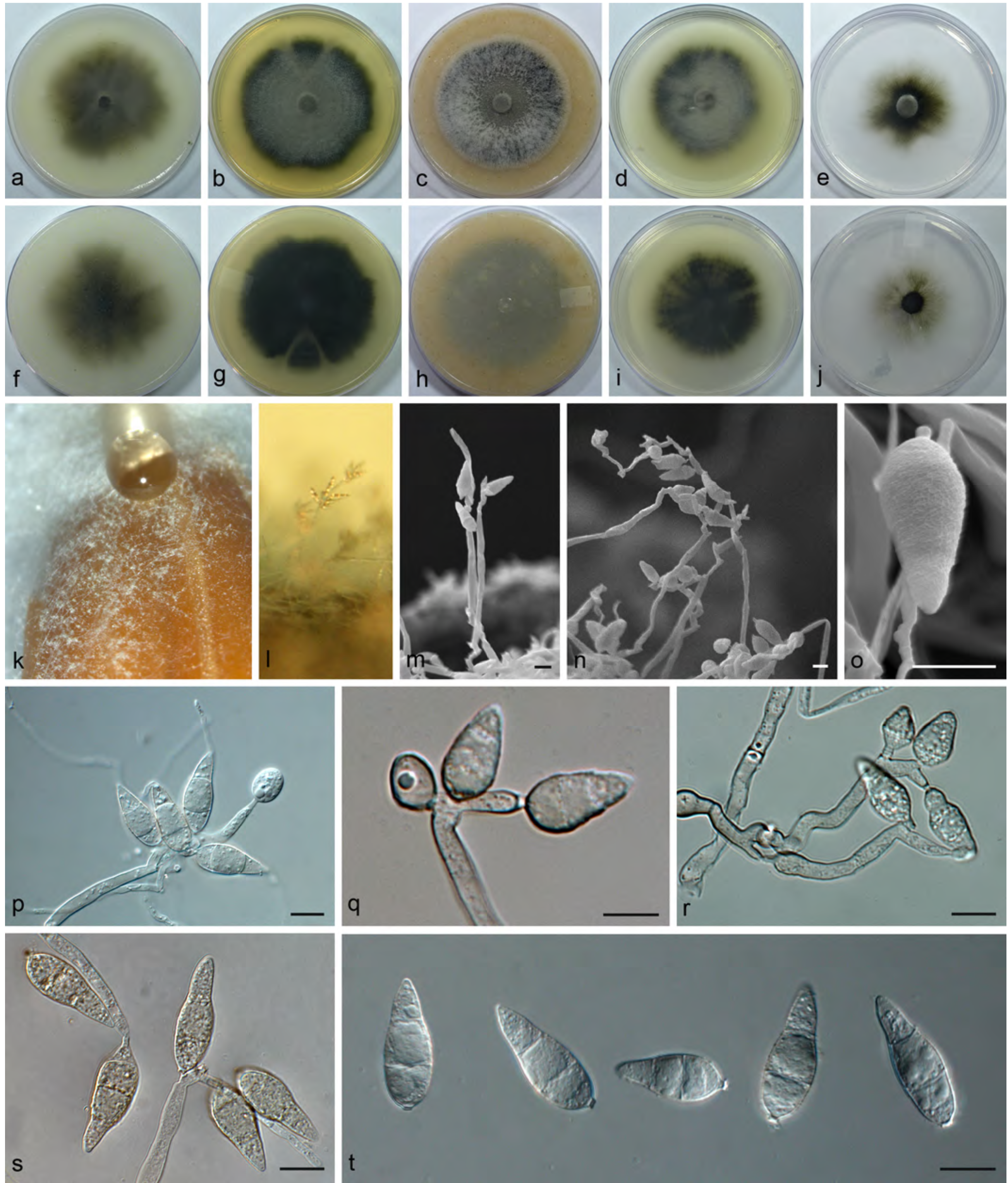
*Pyricularia oryzae* pathotype *Oryza;* a—j: cultures of isolate 10880 grown for 7 d at 12 h photoperiod and 25°C in CMA (a, f), MEA (b, g), OA (c, h), PDA (d, i), and SNA (e, j) media; k—l, sporulation on SNA on sterile barley seeds; m—o, scanning electron micrographs of conidiophores and conidia; p—t, bright field microscopy images of conidiophores and conidia. – Scale bars = 10 μm.

**Fig. 6.**
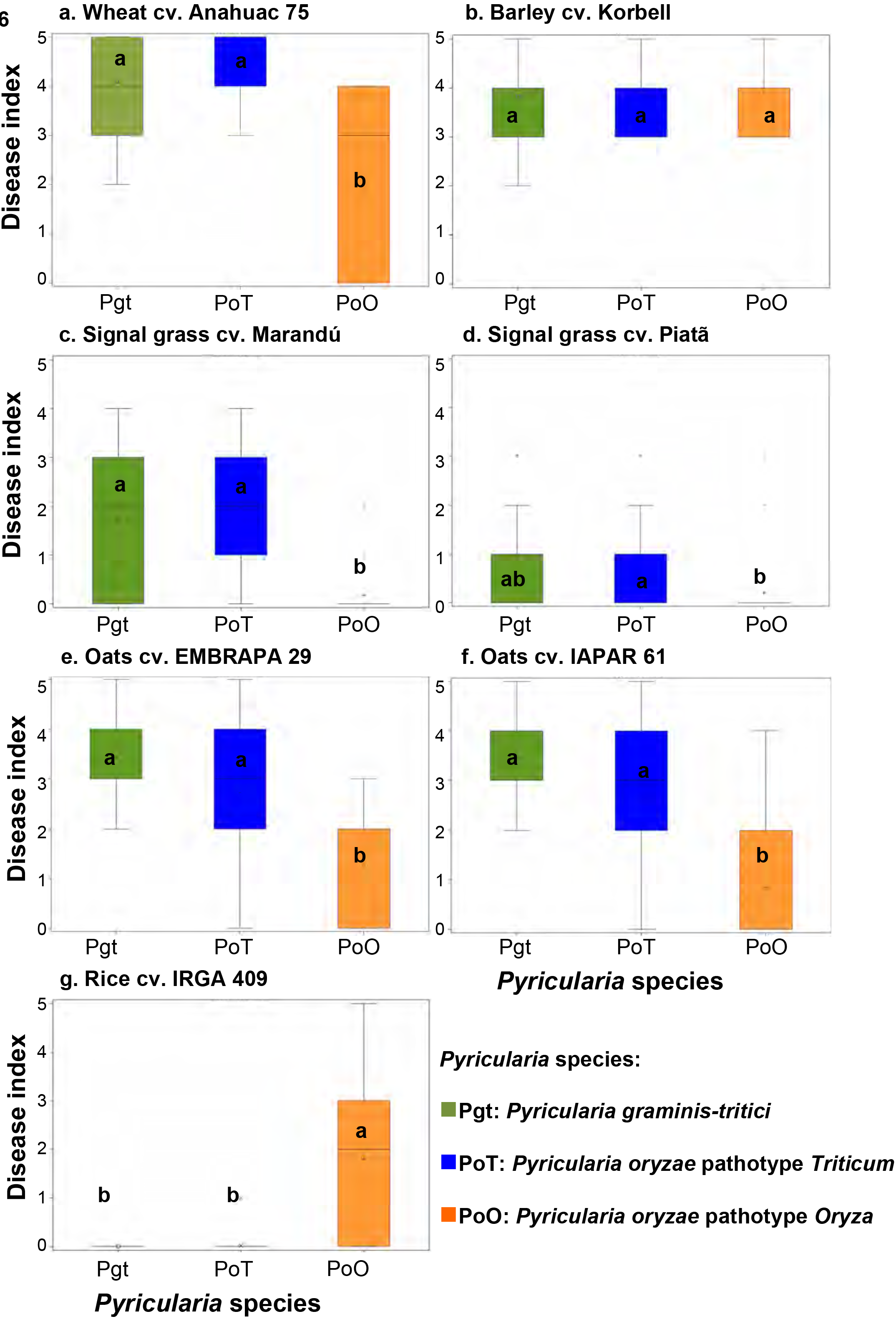
Boxplot distribution of leaf blast severity of seedlings of five poaceous hosts as response to inoculations with isolates of *P. graminis-tritici* (Pgt, *N* = 7), *P. oryzae* pathotype *Triticum* (PoT, *N* = 7), and *P. oryzae* pathotype *Oryza* (PoO, *N* = 4). Boxplots represent blast severity as mean disease index assessed 7 d after inoculation using an ordinal scale from 0 to 5, and based on lesion type (Urashima et al. 2005). Disease index means with the same letter are not significantly different according to Dunn's All Pairs for Joint Ranks non-parametric test (P > χ^2^ ≤ 0.05). a, inoculated seedling of wheat (*Triticum aestivum*); b, barley (*Hordeum vulgare*) cv. BRS Korbell; c, signal grass (*Urochloa brizantha*, ex *Brachiaria brizanta*) cv. Marandú; c, signal grass cv. Piatã; e, oats (*Avena sativa*) cv. EMBRAPA 29; f, oats cv. IAPAR 61; g, rice (*Oryza sativa*) cv. IRGA 409.

**Table 6.**
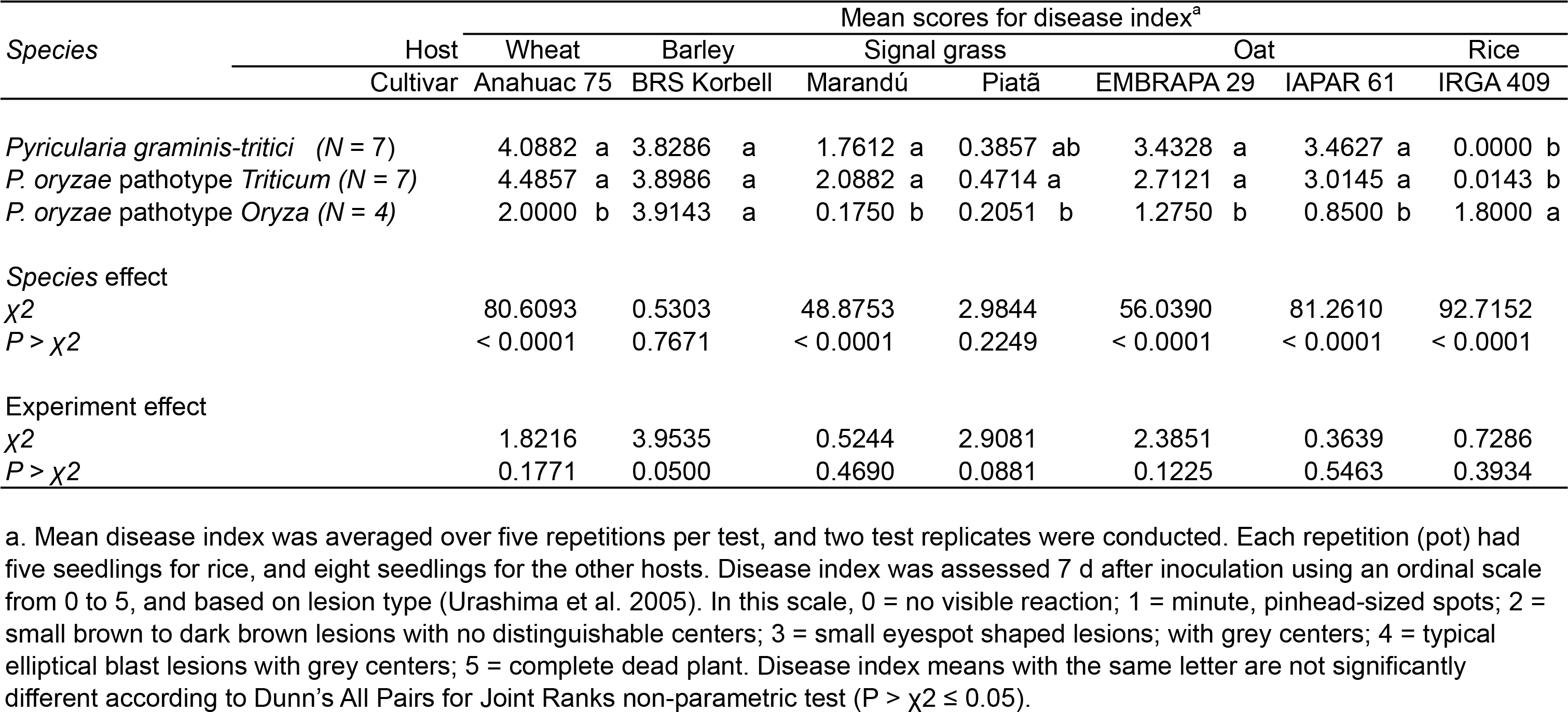
Pathogenicity of isolates of *Pyricularia* spp. on seedlings of five poaceous hosts

Inoculation tests on seedlings of wheat cv. Anahuac 75 showed significant differences among *Pyricularia* species in pathogenicity (mean disease index, DI) (P > χ^2^ < 0.0001). Seedlings were highly susceptible to isolates of PoT and Pgt (DIs of 4.48 and 4.09, respectively). In addition, isolates of PoO caused lesions on wheat seedlings (DI = 2.00); however, conspicuous differences were observed in the levels of virulence of isolates of this group. Isolates 8762 and 10659 sporadically produced lesions that ranged from minute, pinhead-sized spots (type 1 lesion) to small eyespot shaped lesions with grey centres (type 3 lesions). On the other hand, isolates 678 and 10880 consistently produced typical elliptical blast lesions with grey centres (type 4 lesions) (Fig. 6a, 8a).

Seedlings of barley cv. BRS Korbell did not show significant differences in their susceptible response to the inoculated *Pyricularia* species (P > χ^2^ = 0.7671). All species were highly virulent on this host (DIs ≥ 3.82), showing that barley is very susceptible to both wheat and rice blast pathogens (Fig. 6b, 8b).

Inoculations on signal grass seedlings showed that cv. Marandú was more susceptible to *Pyricularia* species than cv. Piatã. On cv. Marandú, PoT (DI = 2.08) showed the highest level of virulence, but it was not significantly different and Pgt (DI = 1.76). PoO was not pathogenic on this cultivar (DI = 0.18). None of the species were pathogenic on signal grass cv. Piatã (DIs ranged from 0.21 to 0.47, and were not significantly different at *P* > χ^2^ = 0.2249) (Fig. 6c, d, 8c).

Inoculation tests on oats showed similar seedling reactions for cvs. EMBRAPA 29 and IAPAR 61. Both Pgt and PoT had similar, high average levels of aggressiveness with DIs > 2.71 for cv. EMBRAPA 29 and DI > 3.01 for cv. IAPAR 61. Furthermore, significant differences in the level of aggressiveness of individual isolates of these species were observed. The most aggressive isolates on oats cv. EMBRAPA 29 were 12.0.534i (Pgt), 12.1.169 and 12.1.119 (both PoT), and the least aggressive isolates were 12.0.607i (Pgt), 12.1.032i and 12.1.291 (both PoT). Likewise, on cv. IAPAR 61 the most aggressive isolates were 12.0.607i (Pgt), 12.1.158 and 12.1.119 (both PoT), and the least aggressive isolates were 12.0.642i (Pgt), 12.0.009i and 12.1.291 (both PoT). Isolates of PoO showed the lowest level of aggressiveness on oats (DI = 1.28 on cv. EMBRAPA 29, and 0.85 on cv. IAPAR 61), significantly lower *P* > χ^2^ < 0.0001) compared to PoT and Pgt. Differences in virulence among isolates of PoO were significant only on cv. IAPAR 61, on which isolate 10659 was the most aggressive while isolate 8762 was not pathogenic (Fig. 6e, f, 8d).

Inoculation tests on rice seedlings showed generally low levels of disease severity. On cultivar IRGA 409, PoO was pathogenic with a mean DI = 1.80 which was significantly different from the DI of the other two species (*P* > χ^2^ < 0.0001). Pgt and PoT were not pathogenic on rice (DI = 0.00 and DI = 0.01, respectively). PoO isolates showed a wide range of aggressiveness. Whereas isolates 8762 and 10880 consistently produced small eyespot-shaped lesions with grey centres (type 3 lesions) and sporadically typical elliptical blast lesions (type 4 lesions), isolate 678 produced small dark brown lesions with no distinguishable centres (type 2 lesions) and isolate 10659 sporadically produced type 2 lesions or no lesions at all on cv. IRGA 409 (Fig. 6h, 8e). This variation in virulence among the isolates is consistent with race-cultivar interactions.

A significant experiment effect was observed in the wheat head inoculation tests (*P* = 0.02). Therefore, statistical analyses of the two test replicates were conducted independently (Table 7, Fig. 7, 8f). The mean disease indexes obtained for PoT and PoO were higher in the second experiment; nevertheless, results from both experiments were congruent. All species tested were pathogenic on heads of wheat cv. Anahuac 75 and significant differences were found in their levels of aggressiveness (*P* < 0.0001 for both experiment 1, and experiment 2). Pgt was the most aggressive species, followed by PoT (Table 7). Isolates of PoO were able to infect wheat heads, but the disease did not progress to more than 10% of the head of cv. Anahuac 75. However, similar to the seedling inoculation tests, PoO isolate 10880 was very aggressive on wheat heads, infecting 20-60% of the inoculated heads (mean DI = 33.39%; Fig. 7, 8f).

**Fig. 7.**
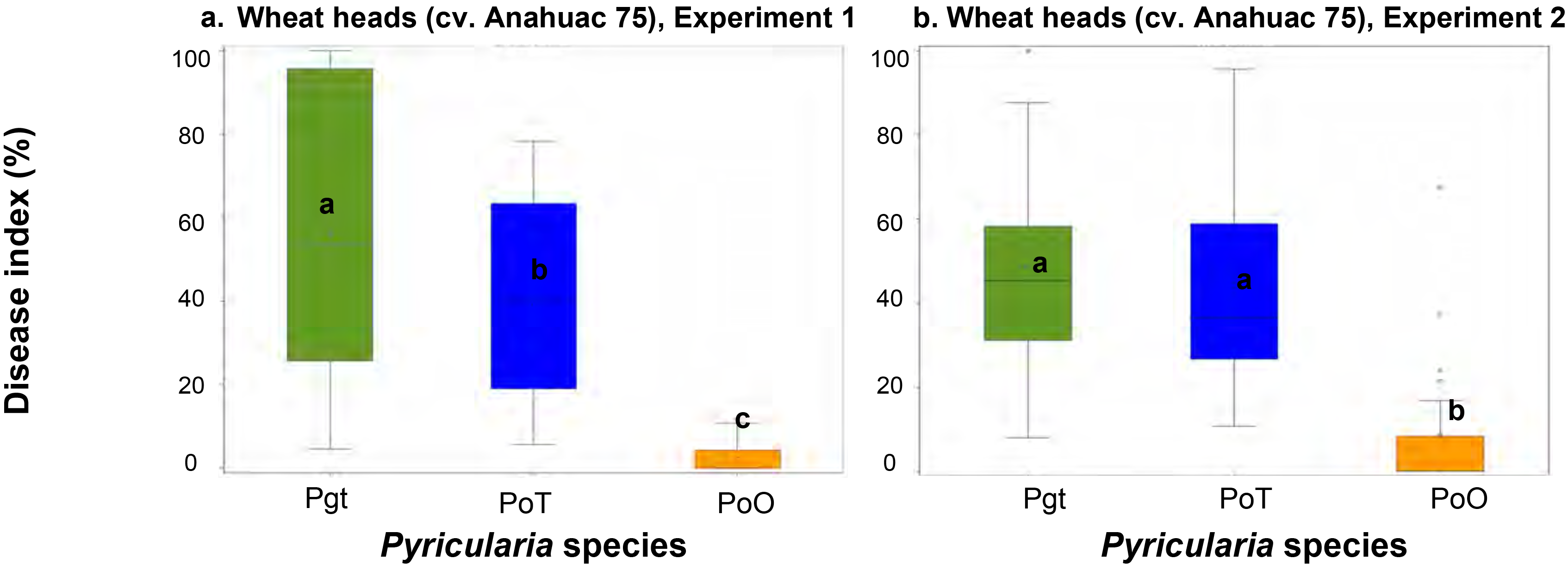
Boxplot distribution of blast severity observed on heads of wheat (*Triticum aestivum*) cv. Anahuac after inoculations with isolates of *P. graminis-tritici* (Pgt, *N* = 7), *P. oryzae* pathotype *Triticum* (PoT, *N* = 7), and *P. oryzae* pathotype *Oryza* (PoO, *N* = 4). Heads were not detached from the plant. Boxplots represent blast severity as mean disease index assessed 7 d after inoculation as percentage wheat head affected by blast using Assess v. 2.0 Image Analysis software. Head tissue was considered diseased when was chlorotic and/or covered in pathogen spores. The test was conducted twice, and replicates (experiment 1 and 2) were analysed independently (a,b). Disease index means with the same letter are not significantly different according to Fisher's protected Least Significant Difference test at P ≤ 0.05.

**Fig. 8.**
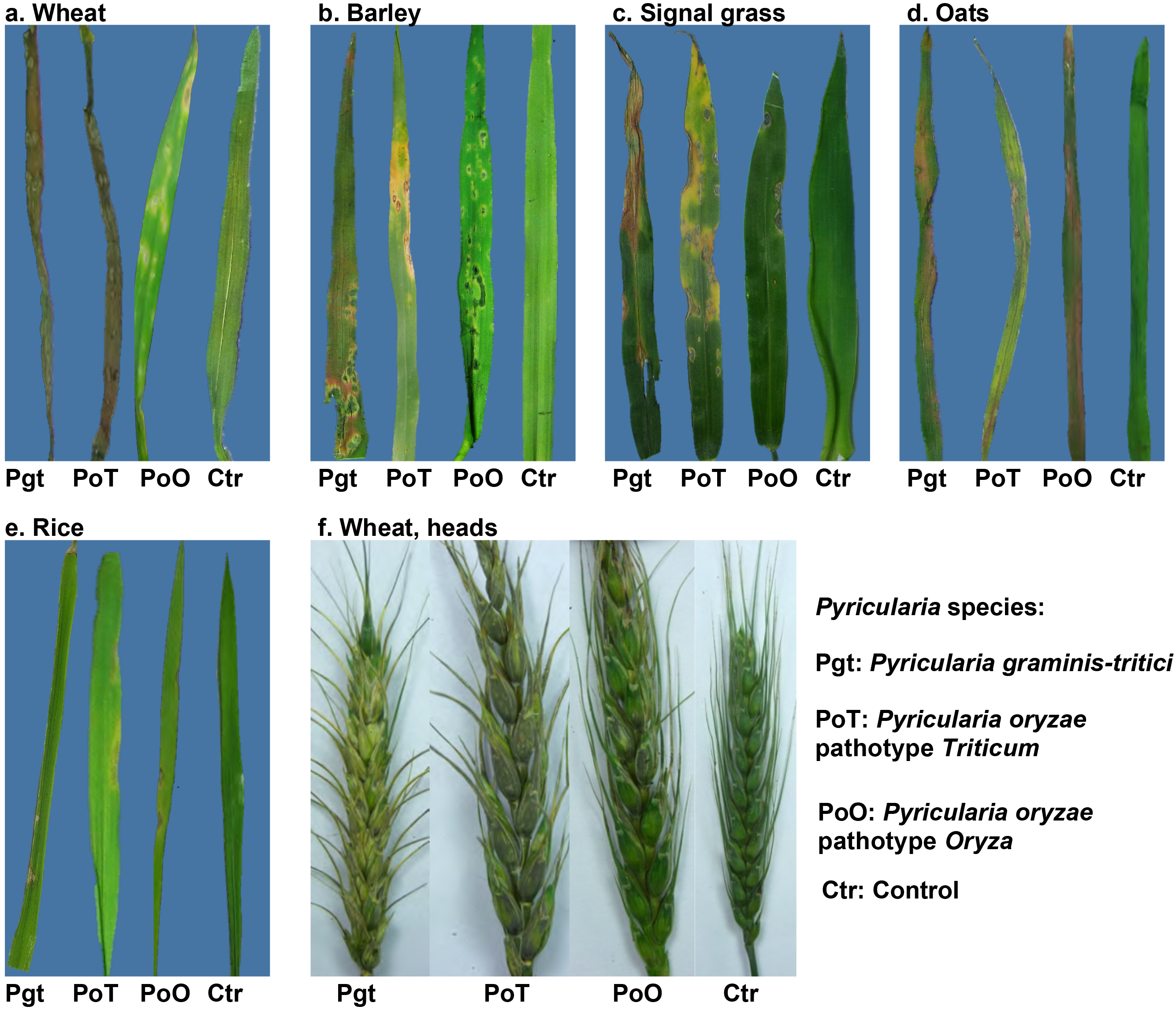
Blast symptoms on leaves and heads of poaceous host after inoculation with *Pyricularia* species. Inoculated hosts: a and f, wheat (*Triticum aestivum*); b, Barley (*Hordeum vulgare*); c, signal grass (*Urochloa brizantha*, ex *Brachiaria brizantha*); d, oats (*Avena sativa*); e, rice (*Oryza sativa). Pyricularia* species: *Pyricularia graminis-tritici* (Pgt), *P. oryzae* pathotype *Triticum* (PoT), and *P. oryzae* pathotype *Oryza* (PoO). Control plants (C) were inoculated with sterile deionized water amended with Tween 80 (2 drops/L). Plants were assessed for disease symptoms 7 d after inoculation.

**Table 7.** Pathogenicity of isolates of *Pyricularia* spp. on non-detached heads of wheat (*Triticum aestivum*) cv. Anahuac 75

| Species, clade | Disease index (% head affected area)<sup>a</sup> |  |  |  |
| --- | --- | --- | --- | --- |
|  | Experiment 1 |  | Experiment 2 |  |
|  | Least Mean Square | Standard Error | Least Mean Square | Standard Error |
| *Pyricularia graminis-tritici* (N = 7) | 57.0364 a | 1.6566 | 47.9202 a | 2.3065 |
| *P. oryzae* pathotype *Triticum* (N = 7) | 39.7740 b | 1.6996 | 43.6509 a | 2.3065 |
| *P. oryzae* pathotype *Oryza* (N = 4) | 2.1330 c | 2.1241 | 8.3485 b | 2.8691 |
| Species effect |  |  |  |  |
| *F* | 209.0400 |  | 65.2000 |  |
| *P* | < 0.0001 |  | < 0.0001 |  |
| LSD | 5.1229 |  | 7.0157 |  |

<sup>a</sup>. Disease index was calculated as the percentage of the wheat head affected by blast using Assess v. 2.0 Image Analysis software. Head tissue was considered diseased when was chlorotic and/or covered in pathogen spores. Disease was assessed 7 d after inoculation. Mean disease index was averaged over five repetitions (wheat heads) for each test replicate. The inoculation experiment was conducted twice, and replicates were analyzed independently due to significant species\*experiment replicate interaction ( $P = 0.0170$ ). Disease index means with the same letter are not significantly different according to Fisher's protected Least Significant Difference (LSD) test at $P \leq 0.05$ .

## TAXONOMY

***Pyricularia graminis-tritici*** V.L. Castroagudm, S.I. Moreira, J.L.N. Maciel, B.A. McDonald, P.W. Crous & P.C. Ceresini, *sp. nov.* — MycoBank MB816086; Fig. 3

*Etymology*. Referring to the major association of this fungal species with multiple grasses, and to the most common cultivated species this fungal species infects causing blast, *Triticum aestivum*.

On SNA on sterile barley seeds — *Mycelium* consisting of smooth, hyaline, branched, septate hyphae, 2-3 μm diam. *Conidiophores* solitary, erect, straight or curved, unbranched, 1-5-septate, medium brown, smooth, (14-)125(-255) × (1-) 3.5(-6) μm. Abundant conidiogenesis observed on the top half of the conidiophore. *Conidiogenous cells* 50-80(-170) × 3-5 μm, terminal and intercalary, pale brown, smooth, forming a rachis with sympodial proliferation, with several protruding denticles, 1-2 μm long, 1.5-2 μm diam. *Conidia* solitary, pyriform to obclavate, pale brown, finely verruculose, granular to guttulate, 2-septate, (23-)25-29(-32) × (8-)9(-10) μm; apical cell 10-13 μm height, basal cell 6-9 μm long; frill hilum, protruding, 1—1.5 μm long, 1.5-2 μm diam, unthickened, not darkened; central cell turning dark brown with age. *Chlamydospores* and *microconidia* not observed.

Culture characteristics — Colonies on CMA with moderate dark grey aerial mycelium, irregular margins, reaching up to 6.5 cm diam after 1 wk; reverse dark grey. Colonies on MEA with abundant white aerial mycelium, and pale grey sporulation at the centre; reaching up to 7.6 cm diam after 1 wk; reverse dark grey; sometimes, fewer colonies (5.1 cm diam) with dark grey sporulation at centre and abundant white aerial mycelium at margins. Colonies on OA with dark grey sporulation in concentric circles, with sparse margins, up to 5.8 cm; reverse pale grey; sometimes, larger growth with abundant white aerial mycelium, pale grey at the centre. Colonies on PDA with abundant white aerial mycelium, olivaceous at centre, growth in concentric circles, up to 6.5 cm diam; reverse black in centre with white margins. Colonies on SNA with sparse olivaceous mycelium irregular margins, up to 5.2 cm diam; reverse sparse olivaceous.

*Typus*. Brazil, Goiás, isolated from head of *Triticum aestivum*, 2012, J.L.N. Maciel (**holotype** as metabolically inactive culture URM7380 = CLM 3547 = isolate 12.1.037).

*Specimens examined*. BRAZIL, Goiás, isolated from head of *Triticum aestivum*, 2012, J.L.N. Maciel (URM7380 = CLM 3547, isolate 12.1.037); Mato Grosso do Sul, isolated from leaves of *Avena sativa*, 2012, J.L.N. Maciel (URM7366 = CLM3516, isolate 12.0.345); Mato Grosso do Sul, isolated from leaves of *Echinochloa crusgalli*, 2012, J.L.N. Maciel (URM7381, isolate 12.0.326); Mato Grosso do Sul, isolated from leaves of *Elionorus candidus*, 2012, J.L.N. Maciel (URM7377, isolate 12.0.194); Mato Grosso do Sul, isolated from leaves of *Urochloa brizantha*, 2012, J.L.N. Maciel (URM7367 = CLM3517, isolate 12.0.366); Paraná, isolated from leaves of *Cenchrus equinatus*, 2012, J.L.N. Maciel (URM7378, isolate 12.0.642i); Paraná, isolated from leaves of *Cynodon spp*., 2012, J.L.N. Maciel (URM7375, isolate 12.0.578i); Paraná, isolated from leaves of *Digitaria sanguinalis*, 2012, J.L.N. Maciel (URM7376, isolate 12.0.555i); Paraná, isolated from leaves of *Eleusine indica*, 2012, J.N.L. Maciel (holotype URM7365 = CLM3518, isolate 12.0.534i); Paraná, isolated from leaves of *Rhynchelytrum repens*, 2012, J.L.N. Maciel (URM7384, isolate 12.0.607i); Rio Grande do Sul, isolated from head of *T. aestivum*, 2012, J.L.N. Maciel (URM7387, isolate 12.1.191).

Notes — *Pyricularia graminis-tritici* causes blast disease on *Triticum aestivum, Avena sativa, Hordeum vulgare* and *Urochloa brizantha* but not on *Oryza sativa*.

Based on morphological and cultural comparisons, isolates of *P. graminis-tritici* are indistinguishable from those of *P. oryzae* pathotypes *Oryza* and *Triticum*. However, these taxa are readily distinguished based on their DNA phylogeny, host range and pathogenicity spectra. Sequencing of the *MPG1* gene is a diagnostic tool to distinguish *P. graminis-tritici* from *P. oryzae*.

***Pyricularia oryzae*** Cavara, Fung. Long. Exsicc. 1: no. 49. 1891. = *Magnaporthe oryzae* B.C. Couch, Mycologia 94: 692. 2002.

***Pyricularia oryzae* pathotype *Triticum*** (Kato et al. 2000); Fig. 4 On SNA on sterile barley seeds — *Mycelium* consisting of smooth, hyaline, branched, septate hyphae, 1.5-2 μm diam. *Conidiophores* solitary, erect, straight or curved, unbranched, medium brown, smooth, 60-150 × 4-6 μm, 2-3-septate; base arising from hyphae, not swollen, lacking rhizoids. *Conidiogenous cells* 40-95 × 3-5 μm, integrated, terminal and intercalary, pale brown, smooth, forming a rachis with several protruding denticles, 0.5-1 μm long, 1.5-2 μm diam. *Conidia* solitary, pyriform to obclavate, pale brown, smooth, granular to guttulate, 2-septate, (25-)27-29(-32) × (8-)9(-10) μm; apical cell 10-13 μm long, basal cell 6-9 μm long; hilum truncate, protruding, 1-1.5 μm long, 1.5-2 μm diam, unthickened, not darkened. *Chlamydospores* and *microconidia* not observed (based on isolate CPC 26580 = 12.1.132).

Culture characteristics — On CMA colonies with moderate dark grey aerial mycelium with irregular margins, sometimes with black aerial mycelium with sporulation in concentric circles, or sparse white mycelial colonies, reaching up to 5.9 cm diam after 1 wk; reverse dark grey with brown margins. On MEA, colonies presented different forms: cottony white aerial mycelia within concentric growth rings, sometimes with a grey sporulation at the centre, reaching up to 6.9 cm diam after 1 wk; reverse dark grey. Colonies on OA with grey aerial mycelium and sporulation in concentric circles; sometimes surface mycelia were white or cream, showing concentric growth, up to 7.9 cm diam; reverse dark grey; sometimes, larger growth with abundant white aerial mycelium, pale grey at the centre. PDA colonies exhibited many variations in culture, often with concentric growth: abundant white aerial mycelia and pale grey sporulation at centre; abundant white aerial mycelia; or dark grey mycelia at the bottom, with white aerial mycelia up to 7 cm diam; reverse, concentric growth, black in centre with olivaceous margins. On SNA the colonies with dark green centres with sparse pale brown margins; or pale grey at the centre and sparse pale brown margins; reverse dark green to black at the centre and with pale brown margins.

*Specimens examined*. BRAZIL, Mato Grosso do Sul, isolated from head of *Triticum aestivum*, 2012, J.L.N. Maciel (URM7388, isolate 12.1.132); Mato Grosso do Sul, isolated from head of *T. aestivum*, 2012, J.L.N. Maciel (URM7368 = CLM3521, isolate 12.1.158); Mato Grosso do Sul, isolated from head of *T. aestivum*, 2012, J.L.N. Maciel (URM7386, isolate 12.1.169); Paraná, isolated from head of *T. aestivum*, 2012, J.L.N. Maciel (URM7369 = CLM3522, isolate 12.1.291); Paraná, isolated from leaves of *Urochloa brizantha*, 2012, J.L.N. Maciel (URM7385, isolate 12.0.009i); Rio Grande do Sul, isolated from head of *T. aestivum*, 2012, J.L.N. Maciel (URM7389, isolate 12.1.205).

***Pyricularia oryzae* pathotype *Oryza*** (Kato et al. 2000); Fig. 5 On SNA on sterile barley seeds. — *Mycelium* consisting of smooth, hyaline, branched, septate hyphae, 2-3 μm diam. *Conidiophores* were (70.5-)146.5(-247) × (3.5-)4.5(-5.5) μm, solitary, erect, straight or curved, septate, hyaline, sometimes light brown. Sometimes, the conidiophores branched. Conidiogeonus cells apical and intercalary, sporulating frequently at the apical part, with protruding *denticles* 0.9-1.1 μm long. *Conidia* pyriform to obclavate, narrowed towards the tip, rounded at the base, 2-septate, hyaline to pale olivaceous, (18-)24-28(-32) × (8-)9(-10) μm; *apical cell* 7-14 μm long, *basal cell* 7-12 μm long; *hilum* 1.5-2 μm diam. *Chlamydospores* and *microconidia* not observed.

Culture characteristics — On CMA the predominant colony morphology was the moderate pale grey aerial mycelium with irregular margins reaching up to 5.6 cm diam after 1 wk; reverse dark grey centre and grey edges; fewer colonies with regular margin formed by sparse white aerial mycelia; sometimes, moderate dark grey aerial mycelium with irregular margins; or white aerial mycelium. Colonies on MEA were often pale grey, sporulation in concentric circles, with dark grey margins; sometimes dark grey at the bottom with sparse white aerial mycelia; or white colonies with regular margins, dark grey at the centre, reaching up to 7.6 cm diam after 1wk; reverse dark grey. On OA colonies with dark grey sporulation at centre and regular margins of white aerial mycelia up to 7.3 cm. PDA colonies were variable, with grey growth in concentric circles, sometimes pale grey or olivaceous; in some cases, with regular margins of white mycelia, reaching up to 6.4 cm; reverse dark grey. On SNA colonies with pale green or dark green mycelia, with sparse margins; in rare cases with abundant pale grey aerial mycelia at centre and white mycelia in regular margins, up to 3.1 cm; reverse dark green in centre and olivaceous at the borders.

*Specimens examined*. BRAZIL, Central Brazil, isolated from leaves of *Oryza sativa*, 2013, Unknown (URM7382, isolate 8762); Central Brazil, isolated from leaves of *O. sativa*, 2013, Unknown (URM7370 = CLM3523, isolate 10880); Goiás, isolated from leaves of *O. sativa*, 2006, Unknown (URM7379, isolate 678); Tocantins, isolated from leaves of *O. sativa*, 2007, Unknown (URM7383, isolate 704).

## DISCUSSION

We conducted comprehensive phylogenetic, morphological and pathogenicity analyses to characterise *Pyricularia* isolates associated with the blast disease on rice, wheat and other poaceous hosts from the Brazilian agro-ecosystem. Urashima, Igarashi & Kato (1993) demonstrated that the blast pathogens infecting wheat and rice were distinct. These authors also reported that isolates recovered from wheat did not infect rice and that most isolates recovered from rice did not infect wheat, except for a few isolates capable of producing small leaf lesions. Although Urashima & Kato (1998), and several follow-up studies demonstrated that the wheat and rice pathogens were phenotypically and genetically different, they have been treated as subgroups of the same species: *Pyricularia oryzae* (Urashima & Kato 1998, Kato et al. 2000, Murakami et al. 2000, Couch & Kohn 2002, Farman 2002, Klaubauf et al. 2014, Chiapello et al. 2015).

The results of our phylogenetic analyses indicate that wheat blast is caused by *Pyricularia* strains assigned to Clade 2, previously described as *P. oryzae* pathotype *Triticum*, and to Clade 3 (Fig. 1, Table 5). Here, we propose that Clade 3 is distinct from *P. oryzae* and represents a new species, *Pyricularia graminis-tritici* (Pgt).

We confirmed that the two host-associated clades *P. oryzae* pathotype *Triticum* and *P. oryzae* pathotype *Oryza* correspond to different pathotypes. This distinction is supported by the combined phylogenetic reconstruction that clearly separates the two taxa. Interestingly, the combined tree (Fig. 2) does not suggest that PoO and PoT are sister taxa. Instead, PoT forms a sister group with Pgt that includes all isolates collected from wheat and other poaceous hosts. This combined group is the sister group to the rice-associated PoO. However, we think that this pattern should be interpreted with caution as explained below.

Among the *Pyricularia* species examined in this study, non-fixed polymorphic sites and phylogenetically informative sites were found in nine of the ten loci examined (locus *BAC6* was monomorphic). Fixed nucleotide differences that are diagnostic for the three taxa were located in four loci: *βT-1, CH7-BAC9, EF-1α* and *MPG1*. Among these, *MPG1* was the most diagnostic locus with 15 fixed differences. Hence, sequencing the *MPG1* locus could provide a simple and informative tool to establish the identity of *Pyricularia* isolates at the species level.

Figure 2 shows the phylogenetic tree reconstructed for *MPG1* using the same settings as described for the combined tree. Significant differences in tree topology are visible compared to the combined tree. Variation at the *MPG1* locus can distinguish Pgt and PoO with high confidence. However, this analysis splits PoT into two sub-clades. Furthermore, PoO and PoT now join together to form the sister-group, as opposed to Pgt. The observation that single loci can produce different phylogenetic patterns has been referred to as “phylogenetic incongruence”. The concept of genealogical concordance of different sequence loci (genealogical concordance phylogenetic species recognition, GCPSR) was proposed as a possible solution for phylogenetic species recognition (Taylor et al. 2000, Dettman et al. 2003). In the GCPSR approach, concordant grouping of species based on several sequences is regarded as evidence for restricted exchange of genetic material and, thus, for the reproductive isolation of taxonomic units, indicating speciation.

However, in an extensive analysis Grünig et al. (2007) showed that this combined phylogenetic approach also has its limits. The authors concluded that in ambiguous cases (such as cryptic species complexes) phylogenetic approaches should be complemented with population genetic analyses that more easily detect reproductive isolation between taxa. Until additional evidence emerges, likely based on comparative population genomics analyses that include entire genome sequences, we suggest a conservative interpretation and propose to maintain the pathotype-based denomination system of *P. oryzae* pathotype *Oryza and P. oryzea Triticum* (Kato et al. 2000), recognizing that PoT and Pgt may eventually be fused into a single, highly diverse species.

Under our experimental conditions, *P. graminis-tritici* and *P. oryzae* pathotypes *Oryza* and *Triticum* did not present consistent cultural or morphological differences. However, distinctive pathogenicity spectra were observed. *Pyricularia graminis-tritici* and *P. oryzae* pathotypes *Triticum* and *Oryza* caused blast symptoms on wheat, barley, and oats with different levels of aggressiveness. These findings agree with Urashima's pioneering observation that two different pyricularia-like pathogens caused wheat blast disease in Brazil (Urashima et al. 2005). Furthermore, our results confirmed that isolates of *P. oryzae* pathotype *Oryza* can cause blast on seedlings and heads of wheat under greenhouse conditions that favour infection, as previously reported (Urashima et al. 1993, Urashima & Kato 1998). An important question that remains to be answered is whether compatible interactions also occur under natural field conditions. Our observation that none of the wheat-derived isolates was genetically assigned to PoO suggests that PoO infections on wheat are very rare or absent under natural field conditions.

In conclusion, our study suggests that blast disease on wheat and other *Poaceae* in Brazil represents a disease complex caused by more than one species of *Pyricularia*. A recent population genomics analysis performed by D. Croll showed that the Bangladeshi wheat blast strains responsible for the 2016 outbreak were closely related to strains of *Pyricularia graminis-tritici* collected in Brazilian wheat fields (Callaway 2016). Given these findings, recognising and properly naming the causal agents of wheat blast will not only increase our understanding of the biology and epidemiology of the disease, but will also enable the establishment of proper quarantine regulations to limit the spread of these pathogens into disease-free areas that grow susceptible wheat cultivars, including Asia, Europe, and North America (McTaggart et al. 2016).

## Acknowledgements

This work was funded by FAPESP (São Paulo Research Foundation, Brazil) research grants to P. C. Ceresini (2013/10655-4 and 2015/10453-8), EMBRAPA/Monsanto research grant (Macroprogram II) to J.L.N. Maciel, and research grants from FINEP (Funding Authority for Studies and Projects, Brazil) and FAPEMIG (Minas Gerais Research Foundation, Brazil) to E. Alves (CAG-APQ-01975-5). P. C. Ceresini and E. Alves were supported by research fellowships from Brazilian National Council for Scientific and Technological Development – CNPq (Pq-2 307361/2012-8 and 307295/2015-0). S.I. Moreira was supported by Doctorate research fellowship from CAPES (Higher Education Personnel Improvement Coordination, Brazil). V. L. Castroagudin was supported by Post-Doctorate research fellowships from CNPq (PDJ 150490/2013-5, from 2012-2014), and FAPESP/CAPES (PDJ 2014/25904-2, from 2015-2016). We thank CAPES for sponsoring the establishment of the “Centro de Diversidade Genética no Agroecossistema” (Pro-equipamentos 775202/2012).

## REFERENCES

Anjos JRND, Silva DBD, Charchar MJD,et al. 1996. Ocurrence of blast fungus (Pyricularia grisea) on wheat and rye in the savanna region of Central Brazil. Pesquisa Agropecuária Brasileira 31: 79–82.

Bozzola JJ, Russell LD. 1999. Electron microscopy: principles and techniques for biologists. Boston: Jones and Bartlett Publishers. p. 670.

Callaway E. 2016. Devastating wheat fungus appears in Asia for first time. Nature 532: 421–422.

Carbone I, Kohn LM. 1999. A method for designing primer sets for speciation studies in filamentous ascomycetes. Mycologia 91: 553–556.

Castroagudm VL, Ceresini PC, Oliveira SC, et al. 2015. Resistance to Qol fungicides is widespread in Brazilian populations of the wheat blast pathogen Magnaporthe oryzae. Phytopathology 104: 284–294.

Chiapello H, Mallet L, Guérin C, et al. 2015. Deciphering genome content and evolutionery relationships of isolates from the fungus Magnaporthe oryzae attacking different hosts plants. Gen Biol Evol 7: 2896–2912.

Choi J, Park S-Y, Kim B-R, et al. 2013. Comparative analysis of pathogenicity and phylogenetic relationship in Magnaporthe grisea species complex. PLoS ONE: 8(2): e57196. doi:57110.51371/journal.pone.0057196.

Couch BC, Fudal I, Lebrun MH, et al. 2005. Origins of host-specific populations of the blast pathogen Magnaporthe oryzae in crop domestication with subsequent expansion of pandemic clones on rice and weeds of rice. Genetics 170: 613–630.

Couch BC, Kohn LM. 2002. A multilocus gene genealogy concordant with host preference indicates segregation of a new species, Magnaporthe oryzae, from M. grisea. Mycologia 94: 683–693.

Crous PW, Verkley GJM, Groenwald JZ, et al. 2009. Fungal Biodiversity. Utrecht, The Netherlands: CBS-KNAW Fungal Biodiversity Centre. p. 1–269.

Cruz MFA, Rios JA, Araujo L, et al. 2016. Infeccion process of Pyricularia oryzae on leaves of wheat seedling. Tropical Plant Pathology DOI 10.1007/s40858-016-0068-6.

Darriba D, Taboada GL, Doallo R, et al. 2012. jModelTest 2: more models, new heuristics and parallel computing. Nature Methods 9: 772.

Dettman JR, Jacobson DJ, Turner E, et al. 2003. Reproductive isolation and phylogenetic divergence in Neurospora: comparing methods of species recognition in a model eukaryote. Evolution 57: 2721–2741.

Drummond AJ, Suchard MA, Xie D, et al. 2012. Bayesian phylogenetics with BEAUti and the BEAST 1.7. Molecular Biology and Evolution 29: 1969–1973.

Duveiller E, Hodson D, Tiedmann A. 2010. Wheat blast caused by Magnaporthe grisea: a reality and new challenge for wheat research. International Wheat Conference, 8: 247–248.

Duveiller E, Singh RP, Nicol JM. 2007. The challenges of maintaining wheat productivity: pests, diseases, and potential epidemics. Euphytica 157: 417–430.

Farman ML. 2002. Pyricularia grisea isolatescausing gray leaf spot on perennial ryegrass (Lolium perenne) in the United States: relationship to P. grisea isolates from other host plants. Phytopathology 92: 245–254.

Grünig CR, Brunner PC, Duò A, et al. 2007. Suitability of methods for species recognition in the Phialocephala fortinii-Acephala applanata species complex using DNA analysis. Fungal Genetins and Biology 44: 773–788.

Hamer JE. 1991. Molecular probes for rice blast disease. Science 252: 632–633.

Hirata K, Kusba M, Chuma I, et al. 2007. Speciation in Pyricularia inferred from multilocus phylogenetic analysis. Mycological Research 111: 799–808.

Igarashi S, Utimada CM, Igarashi LC, et al. 1986. Pyricularia em trigo. 1. Ocorrência de Pyricularia spp. no estado do Paraná. Fitopatologia Brasileira 11: 351–352.

Kato H, Yamamoto M, Yamaguchi-Ozaki T, et al. 2000. Pathogenicity, mating ability and DNA restriction fragment length polymorphisms of Pyricularia populations isolated from Gramineae, Bambusideae and Zingiberaceae plants. Journal of General Plant Pathology 66: 30–47.

Klaubauf S, Tharreau D, Fournier E, et al. 2014. Resolving the polyphyletic nature of Pyricularia (Pyriculariaceae). Studies in Mycology 79: 85–120.

Kohli MM, Mehta YR, Guzman E, et al. 2011. Pyricularia blast – a threat to wheat cultivation. Czech Journal of Genetics and Plant Breeding 47: S130–S134.

Librado P, Rozas J. 2009. DnaSP v5: A software for comprehensive analysis of DNA polymorphism data. Bioinformatics 25: 1451–1452.

Lima MIP, Minella E. 2003. Ocurrence of head blast in barley. Fitopatologia Brasileira 28: 207.

Luo J, Zhang N. 2013. Magnaporthiopsis, a new genus in Magnaporthaceae (Ascomycota). Mycologia 105: 1019–1029.

Maciel JLN. 2011. Magnaporthe oryzae, the blast pathogen: current status and options for its control. Plant Science Reviews 2011: 233–240.

Maciel JLN, Ceresini PC, Castroagudin VL, et al. 2014. Population structure and pathotype diversity of the wheat blast pathogen Magnaporthe oryzae 25 years after its emergence in Brazil. Phytopathology 104: 95–107.

McTaggart AR, van der Nest MA, Steenkamp ET, et al. 2016. Fungal genomics challenges the dogma of name-based biosecurity. PLoS Pathogens 12: e1005475. doi:1005410.1001371/journal.ppat.1005475.

Murakami J, Tomita R, Kataoka T, et al. 2003. Analysis of host species specificity of Magnaporthe grisea toward foxtail millet using a genetic cross between isolates from wheat and foxtail millet. Phytopathology 93: 42–45.

Murakami J, Tosa Y, Kataoka T, et al. 2000. Analysis of host species specificity of Magnaporthe grisea toward wheat using a genetic cross between isolates from wheat and foxtail millet. Phytopathology 90: 1060–1067.

Murata N, Aoki T, Kusaba M, et al. 2014. Various species of Pyricularia constitute a robust clade distinct from Magnaporthe salvinii and its relatives in Magnaporthacea. Journal of General Plant Pathology 80: 66–72.

Rambaut A, Suchard MA, Xie D, et al. 2014. Tracer v1.6, Available from http://beast.bio.ed.ac.uk/Tracer.

Silue DJ, Nottéghem JL, Tharreau D. 1992. Evidence for a gene-for-gene relationship in the Oryza sativa-Magnaporthe grisea pathosystem. Phytopathology 82: 577–580.

Silva CP, Nomura E, Freitas EG, et al. 2009. Efficacy of alternative treatments in the control of Pyricularia grisea on wheat seeds. Tropical Plant Pathology 34: 127–131.

Takabayashi N, Tosa Y, Oh HS, et al. 2002. A gene-for-gene relationship underlying the species-specific parasitism of Avena/Triticum isolates of Magnaporthe grisea on wheat cultivars. Phytopathology 92: 1182–1188.

Taylor JW, Jacobson DJ, Kroken S, et al. 2000. Phylogenetic species recognition and species concepts in fungi. Fungal Genetics and Biology 31: 21–32.

Tosa Y, Chuma I. 2014. Classification and parasitic specialization of blast fungi. Journal of General Plant Pathology 80: 202–209.

Tosa Y, Hirata K, Tamba H, et al. 2004. Genetic constitution and pathogenicity of Lolium isolates of Magnaporthe oryzae in comparison with host species-specific pathotypes of the blast fungus. Phytopathology 94: 454–462.

Tosa Y, Tamba H, Tanaka K, et al. 2006. Genetic analysis of host species specificity of Magnaporthe oryzae isolates from rice and wheat. Phytopathology 96: 480–484.

Urashima AS, Galbieri R, Stabili A. 2005. DNA fingerprinting and sexual characterization reveled two distinc populations of Magnaporthe grisea in wheat blast in Brazil. Czech Journal of Genetics and Plant Breeding 41: 238–245.

Urashima AS, Hashimoto Y, Don LD, et al. 1999. Molecular analysis of the wheat blast population in Brazil with s homolog of retrotransposon MGR583. Annals of the Phytopathological Society of Japan 65: 429–436.

Urashima AS, Igarashi LC, Kato H. 1993. Host range, mating type, and fertility of Pyricularia grisea from wheat in Brazil. Plant Disease 77: 1211–1216.

Urashima AS, Kato H. 1998. Pathogenic relationship between isolates of Pyricularia grisea of wheat and other hosts at different host developmental stages. Fitopatologia Brasileira 23: 30–35.

Valent B, Chumley FG. 1991. Molecular genetic analysis of the rice blast fungus, Magnaporthe grisea. Annual Review of Phytopathology 29: 443–467.

Valent B, Khang CH. 2010. Recent advances in rice blast effector research. Current Opinion in Plant Biology 13: 434–441.

Verzignassi RS, S. PL, Benchimol RL, et al. 2012. Pyricularia grisea: new pathogen on Brachiaria brizantha cv. Marandú in Para. Summa Phytopathologica 38: 254.

Zadocks JC, Chang TT, Konzak CF. 1974. A decimal code for the growth stages of cereals. Weed Research 14: 415–421.

Zhang N, Zhao S. 2011. A six-gene phylogeny reveals the evolution of mode of infection in the rice blast fungus and allied species. Mycologia 103: 1267–1276.

